# Cytotoxic CNS-associated T cells drive axon degeneration by targeting perturbed myelinating oligodendrocytes in *PLP1* mutant mice

**DOI:** 10.1101/2023.01.10.523231

**Authors:** Tassnim Abdelwahab, David Stadler, Konrad Knöpper, Panagiota Arampatzi, Antoine-Emmanuel Saliba, Wolfgang Kastenmüller, Rudolf Martini, Janos Groh

**Affiliations:** Department of Neurology, Section of Developmental Neurobiology, University Hospital Würzburg, Würzburg, Germany; Institute for Systems Immunology, University of Würzburg, Würzburg, Germany; Core Unit Systems Medicine, University of Würzburg, Würzburg, Germany; Helmholtz Institute for RNA-based Infection Research, Helmholtz-Center for Infection Research, Würzburg, Germany

**Keywords:** axon, myelin, neuroinflammation, neurodegeneration, CD8^+^ T cells, fingolimod

## Abstract

Myelin defects lead to neurological dysfunction in various diseases and in normal aging. Chronic neuroinflammation often contributes to axon-myelin damage in these conditions and can be initiated and/or sustained by perturbed myelinating glia. We have previously shown that distinct mutations in the *PLP1* gene result in neurodegeneration that is largely driven by adaptive immune cells. Here we characterize CD8^+^ CNS-associated T cells in these myelin mutants using single-cell transcriptomics and identify population heterogeneity and disease-associated changes. We demonstrate that early sphingosine-1-phosphate receptor modulation attenuates the recruitment of T cells and neural damage, while later targeting of CNS-associated T cell populations is inefficient and has no effect on neurodegeneration. Applying bone marrow chimerism and utilizing random X chromosome inactivation, we provide evidence that axonal damage is driven by cytotoxic, antigen specific CD8^+^ T cells that target mutant myelinating oligodendrocytes. These findings offer insights into neural-immune interactions and are of translational relevance for neurological conditions associated with myelin defects and neuroinflammation.

## Introduction

The integrity of myelinated axons is essential for the proper function of the mammalian central nervous system (CNS)^1^. Due to their unique properties, myelinated axons are particularly susceptible to injury and their perturbation is a hallmark and early feature of various neurological diseases and of normal aging^2,3^. Recent findings have highlighted the important interplay between neural cells and immune cells in maintaining homeostasis of the axon-myelin unit^4–8^. Disturbances of this interaction can contribute to the initiation and perpetuation of neuroinflammation, demyelination, axonal damage, and neurodegeneration, contributing to functional decline and clinical impairment in distinct neurological diseases such as multiple sclerosis, Alzheimer’s disease, Parkinson’s disease, hereditary diseases, and in normal aging^9,10^.

In these conditions, macroglial (oligodendrocytes, astrocytes) and microglial cells exhibit changes in their gene and protein expression and adopt a chronic pro-inflammatory state^4,7,10,11^. Many of these glial pro-inflammatory changes are related to the communication with adaptive immune cells, including elevated expression of molecules implicated in antigen presentation, T cell receptor stimulation/co-stimulation, and T cell recruitment^8^. Along these lines, there is increasing evidence for a contribution of T lymphocytes and particularly CD8^+^ T cells to many neurological disorders including inflammatory and classical neurodegenerative diseases, often associated with aging^12–14^.

We have previously shown that secondary neuroinflammation acts as an important and targetable amplifier of neural damage in distinct genetically mediated CNS diseases^15^. In mice overexpressing normal or carrying mutant proteolipid protein (*PLPtg* and *PLPmut* mice, respectively), the major myelin protein of the CNS, adaptive immune cells accumulate in the white matter, drive axonopathic and demyelinating alterations, and contribute to functional impairment^16–19^. Pharmacological targeting of innate and adaptive immune reactions can attenuate disease progression in the respective models but has only limited potential to reverse functional impairment^20,21^. This has important implications for progressive forms of multiple sclerosis and leukodystrophies/hereditary spastic paraplegia, which are associated with chronic low-grade neuroinflammation and axon degeneration and can be related to primary oligodendrocyte perturbation^15,22–24^. Moreover, we recently identified commonalities among normal aging and mice with myelin gene defects, underscoring the broad relevance of these processes for frequent aging-related diseases. We showed that in normal aging (wildtype) mice without defined disease, cytotoxic CD8^+^ CNS-associated T cells drive axonal damage and neurodegeneration^25^. Similar as in *PLPtg* mice^19^, this process is dependent on the cytolytic effector protease granzyme B and cognate TCR specificity. Moreover, T cell-driven axon degeneration in aged mice can be aggravated by mimicking infection-related systemic inflammation^25^. Thus, also in aging, perturbation of white matter glial cells results in secondary myelin-related neuroinflammation and contributes to structural and functional decline of myelinated axons. The comparison of these processes between normal aging mice and models with cell type-specific myelin defects might help to clarify some of the involved pathomechanisms and identify targets for intervention.

Both our previous characterization of CD8^+^ T cells in the CNS of adult and aged mice and pharmacological treatment approaches in *PLPmut* mice indicated population heterogeneity and functional diversity of these cells^21,25^. However, the exact composition of CD8^+^ T cell populations and their disease-related changes in the CNS of *PLPmut* mice have not been analyzed. Moreover, their recruitment and maintenance as well as putative pathogenic effector mechanisms and target structures have not been characterized. These issues are of high relevance for strategies to attenuate deterioration of the nervous system by targeting chronic neuroinflammation. Here, we focus on these questions in the context of sphingosine-1-phosphate receptor (S1PR) modulation with fingolimod, an established disease-modifying treatment for multiple sclerosis that sequesters lymphocyte subsets in secondary lymphoid organs^26^. This might offer novel explanations for its limited efficacy in progressive disease forms and lead to refined indications for immunomodulatory therapy in multiple neurological disorders.

## Results

### Transcriptional signatures of CD8^+^ T cells associated with the healthy and myelin mutant CNS

To characterize CD8^+^ CNS-associated T lymphocytes in detail we used single-cell RNA sequencing (scRNA-seq) of CD8^+^ T cells isolated from the brains of adult (12-month-old) wildtype (*Wt*) and *PLP1* mutant (*PLPmut*) mice. Unsupervised clustering of the combined datasets identified nine different clusters (Fig. 1 A), revealing population heterogeneity. Similar as previously observed when comparing cells from the same adult with aged mice^25^, two clusters resembled central memory T (TCM) cells (TCM1 and TCM2; Fig. 1 B). Moreover, we identified one cluster representing effector T (TEFF) cells and one cluster with a prominent interferon response signature (interferon-stimulated T cells (IST)). We previously localized these subsets in the blood, cerebrospinal fluid (CSF), leptomeninges and choroid plexus of adult and aged mice^25^. We also detected the presence of five groups of previously described CD8^+^ CNS-associated T cells (CAT1-5) with distinct transcriptional signatures. CAT1, previously found strongly enriched in white matter of aging brains and expressing inhibitory checkpoint molecules like *Lag3* and *Pdcd1*, was similarly frequent in *Wt* and *PLPmut* mice. This difference in cluster allocation in comparison with aged mice is likely due to the similarity of these cells to CAT3 and CAT4 in *PLPmut* mice. Myelin disease-related changes in CD8^+^ T cells were primarily based on increased numbers of cells representing CAT2 to CAT5 and transcriptional changes within CAT2, e.g., increased expression of *Ly6a* and *Klrc1* (Fig. 1 C-G).

**Figure 1.**
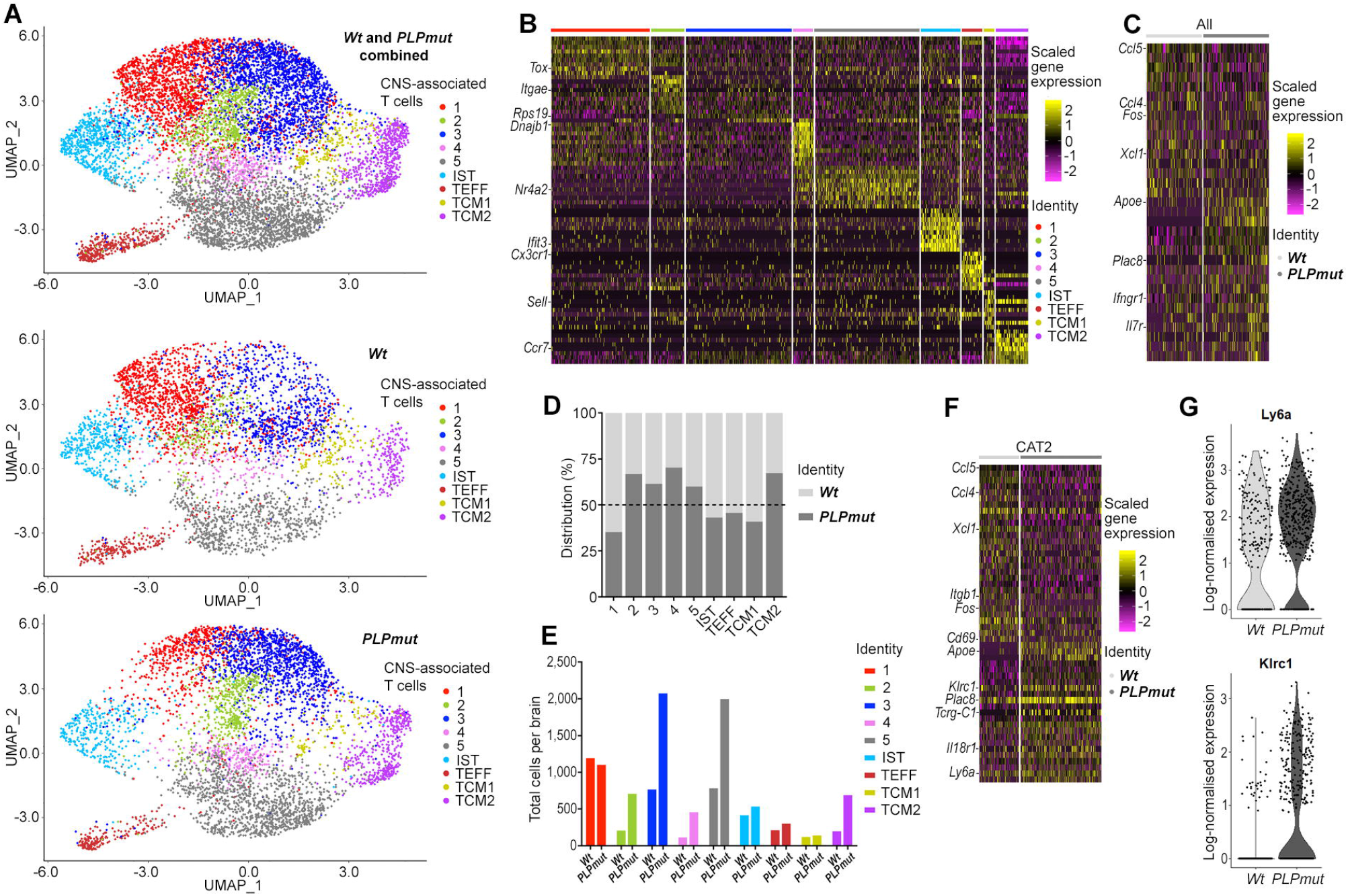
scRNA-seq reveals heterogeneity and activation of CD8^+^ T lymphocytes in the myelin mutant CNS. (**A**) UMAP (uniform manifold approximation and projection) visualization of CD8^+^ T cells pooled and freshly sorted from adult (12-month-old) *Wt* (*n* = 5) and *PLPmut* (*n* = 4) mouse brains and analyzed by scRNA-seq. Combined (top, 9,448 cells) and separate visualization of cells from *Wt* (middle, 4,338 cells) and *PLPmut* (bottom, 5,110 cells) brains are displayed. (**B**) Heatmap of top 10 cluster-specific genes. The color scale is based on a z-score distribution from −2 (purple) to 2 (yellow). Complete lists of cluster-specific marker genes can be found in Table S1. (**C**) Heatmap of differentially expressed genes comparing cells isolated from *Wt* and *PLPmut* brains across all clusters (**D**) contribution of the samples to each cluster is displayed in percent and (**E**) absolute numbers extrapolated to total cells per brain. (**F**) Heatmap of differentially expressed genes between *Wt* and *PLPmut* mice within CAT2 as annotated in panel A. (**G**) Violin plots of *Ly6a* (top) and *Klrc1* (bottom) gene expression in CAT2. CAT, CNS-associated T cells; IST, interferon-stimulated T cells; TEFF, effector T cells; TCM, central memory T cells.

CD8^+^ CNS-associated T cells showed variable expression of gene modules related to effector and memory function and high expression levels for markers of tissue recruitment and residency (Fig. 2 A). Flow cytometry of CD8^+^ T lymphocytes isolated from brains of *Wt* and *PLPmut* mice confirmed the presence of distinct populations and strongly increased numbers of Ly6A/E^+^CD103^+^ cells in *PLPmut* mice (Fig. S1). These correspond to CAT2, which expressed the highest amount of *Gzmb* and *Itgae* (encoding CD103) among CD8^+^ CAT (Table S1). We furthermore validated the heterogenous expression of different identified CD8^+^ T cell subset markers in the white matter of *Wt* and *PLPmut* mice by immunofluorescence (Fig. S2). Again, a disproportional increase of CD8^+^CD103^+^ T cell numbers with an increased expression of Ly6A/E was detectable in the myelin mutants.

**Figure 2.**
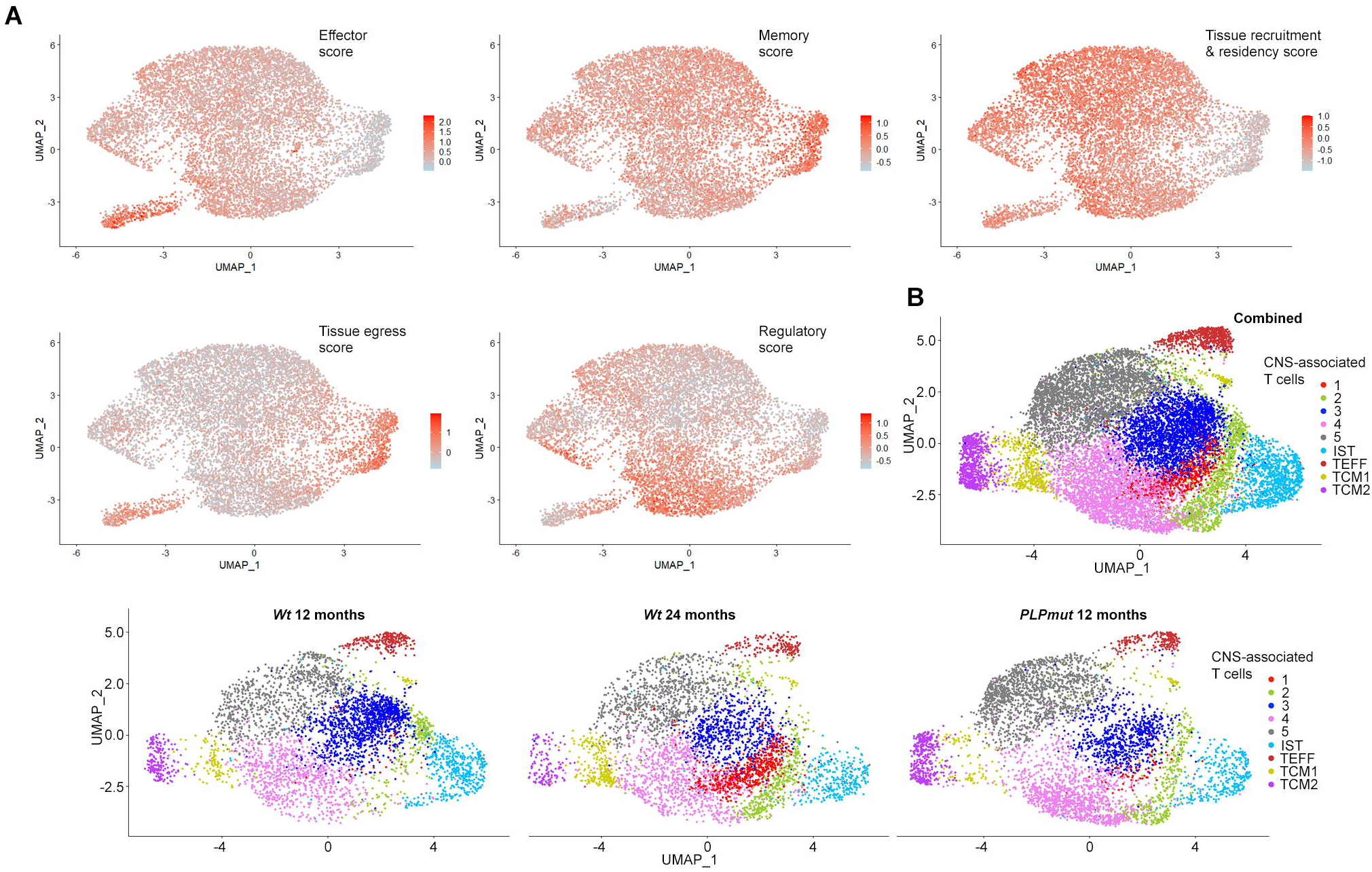
scRNA-seq reveals signatures and commonalities of CD8^+^ CNS-associated T cells in myelin disease and normal aging. (**A**) UMAP visualizations of CD8^+^ T cells from brains of *Wt* and *PLPmut* mice showing the expression of selected module scores (as annotated in Fig. 1 A). Transcript levels are color-coded: lightblue, not expressed; red, expressed. Effector score: *Prf1*, *Gzma*, *Gzmb*, *Fasl*, *Ifng*, *Klrc1*, *Klrg1*, *Cx3cr1*, *Lgals3*, *Ccl5*; Memory score: *Il7r*, *Cd44*, *Ltb*, *Tcf7*, *Cd27*, *Ccr7*, *Sell*, *Fas*; Tissue recruitment & residency score: *Cxcr6*, *Cxcr3*, *Ccr5*, *Gzmk*, *Itgb1*, *Itgb2*, *Cd69*, *Itga1*, *Itgae*, *Tox*, *S100a4*, *S100a6*; Tissue egress score: *Sell*, *Ccr7*, *S1pr1*, *S1pr4*, *S1pr5*, *Klf2*, *Lef1*; Regulatory score: *Nr4a1*, *Nr4a2*, *Pdcd1*, *Il2rb*, *Tnfaip3*, *Ccl4*, *Cxcr4*, *Icos*, *Tgif1*, *Cd28*, *Nfkbid*, *Tnfrsf1b*, *Il10*. (**B**) UMAP visualization of CD8^+^ T cells pooled and freshly sorted from adult (12-month-old) *Wt* (*n* = 5), aged (24-month-old) *Wt* (*n* = 4), and adult *PLPmut* (*n* = 4) mouse brains and analyzed by scRNA-seq. Combined (top, 13,919 cells) and separate visualization of cells from adult *Wt* (middle, 4,338 cells), aged Wt (4,471 cells), and adult *PLPmut* (bottom, 5,110 cells) brains are displayed.

To compare CD8^+^ T cells accumulating in the brains of aged and *PLPmut* mice, we integrated the present and previous datasets of adult and aged (24-month-old) *Wt*, and adult *PLPmut* mice. We confirmed that the originally identified CAT1 population is specifically enriched in aging mice and that allocation of cells from CAT3 and CAT4 to this cluster occurs when comparing adult *Wt* and *PLPmut* mice due to their similar transcriptional profiles (Fig. 2 B). In summary, CD8^+^ CNS-associated T cells are heterogeneous and show commonalities but also differences in genetic myelin disease compared with normal aging. Their transcriptional signatures indicate cytotoxic effector memory function, activation, and long-term tissue residency^27^.

### Distinct S1PR modulation regimens reveal recruitment dynamics and maintenance of CNS-associated T cells in myelin disease

Our unbiased analysis of CD8^+^ CAT indicated that they show relatively strong expression of genes associated with tissue recruitment (e.g., *Cxcr6*, *Cxcr3*, *Gzmk*) and residency (e.g., *Cd69*, *Itga1*, *Itgae*), while marker genes for tissue egress and recirculation (e.g., *S1pr1*, *S1pr3*, *S1pr5*) were mostly confined to TCM and TEFF clusters (Figs. 2 A and 3 A). This might reflect their adaption to maintenance in the brain and has implications for immunomodulatory approaches that antagonize S1PR signaling to target T cell-mediated neuroinflammation. To address this question, we performed pharmacological treatment of *PLPmut* mice using fingolimod (FTY720) in the drinking water and distinct treatment regimens. Based on our previous characterization of chronic T cell recruitment and disease progression in the myelin mutant model from around 2 months of age onwards^16^, we started early and late onset regimens at 4 and 10 months of age, respectively (Fig. 3 B). Fingolimod modulates S1PR signaling to sequester lymphocyte subsets in secondary lymphoid organs^26^, and we confirmed a strong depletion of circulating T cells in *PLPmut* mice by the treatment (Fig. S3 A and B). Lymphopenia was reflected by a significant reduction in relative spleen weight (Fig. S3 B and C).

**Figure 3.**
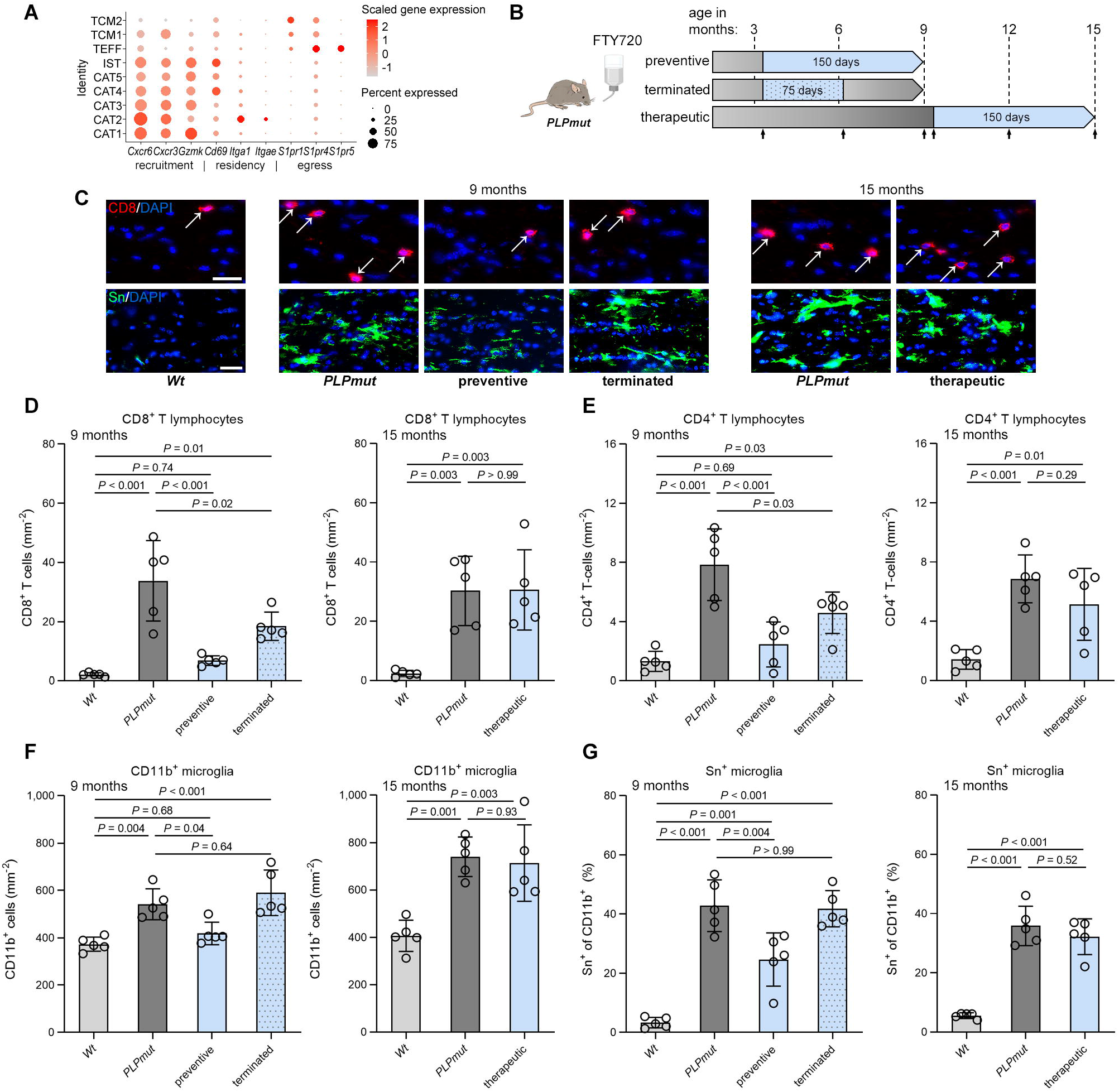
Preventive but not therapeutic sphingosine-1-phoshpate receptor modulation attenuates neuroinflammation in *PLPmut* mice. (**A**) Dot plot expression visualization of selected genes implicated in tissue recruitment, residency, and egress for CD8^+^ T cell subtypes from brains of *Wt* and *PLPmut* mice (as annotated in Fig. 1 A). The color scale is based on a z-score distribution from −1 (lightgrey) to 2 (red). (**B**) Schematic experimental design for distinct treatment regimens using fingolimod (FTY720) in the drinking water. Arrows indicate timepoints for non-invasive analysis by OCT (see Fig. 4). (**C**) Immunofluorescence detection of CD8^+^ T lymphocytes (top; arrows) as well as Sn^+^ activated microglia (bottom) in the optic nerves of *Wt*, *PLPmut* and FTY720-treated *PLPmut* mice using regimens indicated in panel B. Scale bars, 20 μm. (**D**) Quantification of CD8+ T cells, (**E**) CD4+ T cells, (**F**) CD11b+ microglia, and (**G**) activated Sn+ microglia (% of CD11b^+^) in the optic nerves of *Wt*, *PLPmut* and FTY720-treated *PLPmut* mice (*n* = 5 mice per group). Preventive FTY720 treatment attenuates T cell recruitment and microgliosis in *PLPmut* mice which is partially maintained after termination at half time. Therapeutic FTY720 treatment has no effect on ongoing neuroinflammation in *PLPmut* mice. D-G: one-way ANOVA with Tukey’s multiple comparisons test. Data are presented as the mean ± SD. All data represent at least three independent experiments.

Focusing on the optic nerve as a model white matter tract, we found that early onset (preventive) treatment for 150 days attenuated the recruitment of CD8^+^ and CD4^+^ T cells in *PLPmut* mice (Fig. 3 C-E). Termination of the treatment at half-time (after 75 days) resulted in restoration of relative spleen weight, reflecting a re-establishment of circulating lymphocytes (Fig. S3 C). However, numbers of CNS-associated T cells were still lower than in untreated mice, demonstrating a relatively slow recruitment without overshoot after cessation of immunomodulation in *PLPmut* mice (Fig. 3 C-E). Late onset therapeutic treatment of *PLPmut* mice with FTY720 resulted in a similar reduction of spleen weight as preventive treatment (Fig. S3 D) but had no effect on T cell numbers in optic nerves (Fig. 3 C-E). Neuroinflammation in the white matter of *PLPmut* mice is characterized by microglial activation, which orchestrates^20^ but also reacts to adaptive immune reactions^16^ and is reflected by increased expression of sialoadhesin (Sn, Siglec-1, CD169; Fig S4). Parenchymal Sn^+^ microglia in *PLPmut* mice also expressed the established marker for disease-associated microglia CD11c and low but detectable levels of P2RY12, corroborating that they do not represent infiltrated monocyte-derived macrophages or substantial numbers of border-associated macrophages^20^. Preventive treatment with fingolimod inhibited the numerical increase of CD11b^+^ microglia in *PLPmut* mice and attenuated their Sn expression (Fig. 3 C, F, and G). In contrast, microgliosis was restored after termination of treatment or unaffected by therapeutic treatment (Fig. 3 C, F, and G). Taken together, once CD8^+^ T cells become CNS-associated in white matter of *PLPmut* mice, they do not respond to S1PR modulation and sustain chronic neuroinflammation.

### Neuroaxonal degeneration in PLPmut mice correlates with the response of CNS-associated T cells to S1PR modulation

Next, we investigated how FTY720 treatment affects axonal damage and neuron loss in *PLPmut* mice. Neuroinflammation-related degeneration of myelinated axons in these mice is preceded by focal block of axonal transport and formation of spheroids, which can be visualized using antibodies against non-phosphorylated neurofilaments^16^. Corresponding to the response of T cells, preventive treatment attenuated axonal spheroid formation in optic nerves of *PLPmut* mice, while therapeutic treatment had no effect on ongoing axonal damage (Fig. 4 A and B). The slow recruitment of T cells after treatment termination translated into preserved axonal integrity compared with untreated mice. Axon degeneration in *PLPmut* mice culminates in neuron loss and can be monitored using non-invasive longitudinal readout measures of the retinotectal system like optical coherence tomography (OCT)^16^. Preventive and terminated treatment regimens significantly decreased loss of RBPMS^+^ retinal ganglion cells and delayed the selective thinning of the inner retinal layers in OCT (Fig. 4 A and C). In contrast, therapeutic treatment did not slow the progression of neuron loss or retinal thinning.

**Figure 4.**
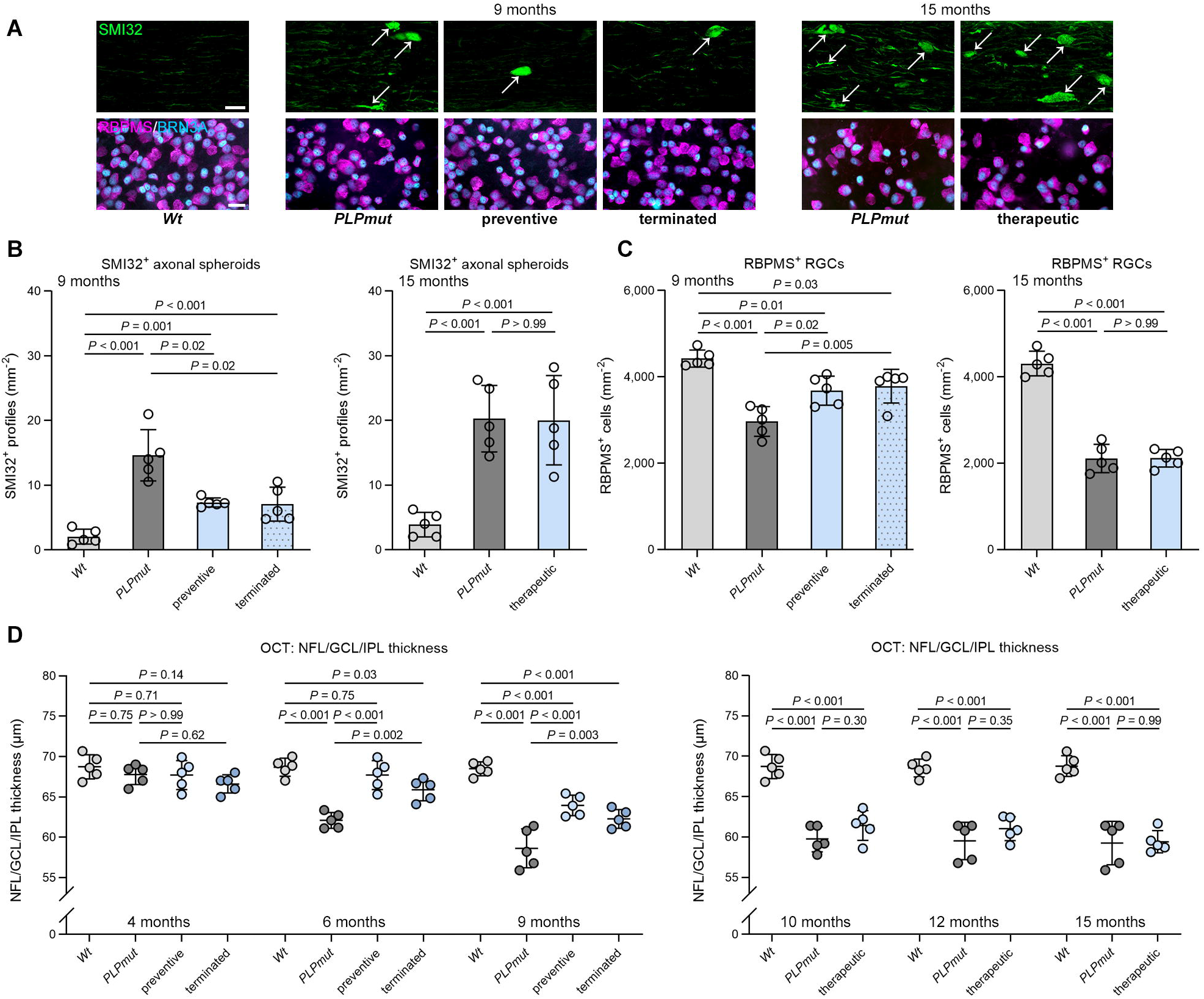
Preventive but not therapeutic FTY720 treatment attenuates neuroaxonal degeneration in *PLPmut* mice. (**A**) Immunofluorescence detection of SMI32^+^ axonal spheroids (top; arrows) in the optic nerves and RBPMS^+^Brn3a^+^ RGCs (bottom) in the retinae of *Wt*, *PLPmut* and FTY720-treated *PLPmut* mice using regimens indicated in Fig. 3 B. Scale bars, 20 μm. (**B**) Quantification of SMI32+ axonal spheroids and (**C**) RGCs in *Wt*, *PLPmut* and FTY720-treated *PLPmut* mice (*n* = 5 mice per group). Preventive FTY720 treatment attenuates axonal damage and neuron loss in *PLPmut* mice which is maintained after termination at half time. Therapeutic FTY720 treatment has no effect on the progression of neurodegeneration in *PLPmut* mice. (**D**) OCT analysis of the innermost retinal composite layer (NFL/GCL/IPL) in peripapillary circle scans. Preventive FTY720 treatment attenuates retinal thinning in *PLPmut* mice which is partially maintained after termination at half time. Therapeutic FTY720 treatment has no effect on retinal thinning in *PLPmut* mice. B, C: one-way ANOVA with Tukey’s multiple comparisons test. D: two-way ANOVA with Tukey’s multiple comparisons test. Data are presented as the mean ± SD. All data represent at least three independent experiments.

We assessed the beneficial effects of preventive FTY720 treatment on white matter integrity in *PLPmut* mice at the ultrastructural level using electron microscopy. The frequency of axons with abnormally thin (g-ratio ≥ 0.85) or no myelin was not significantly different between treated and untreated groups (Fig. 5 A and B). However, corroborating our immunohistochemical observations, axons undergoing swelling and organelle accumulation, or Wallerian-like degeneration were less frequent after fingolimod (Fig. 5 A and C). Thus, neurodegeneration in *PLPmut* mice can be attenuated by early preventive but not by late S1PR modulation, being in line with the effects on CNS-associated T cells.

**Figure 5.**
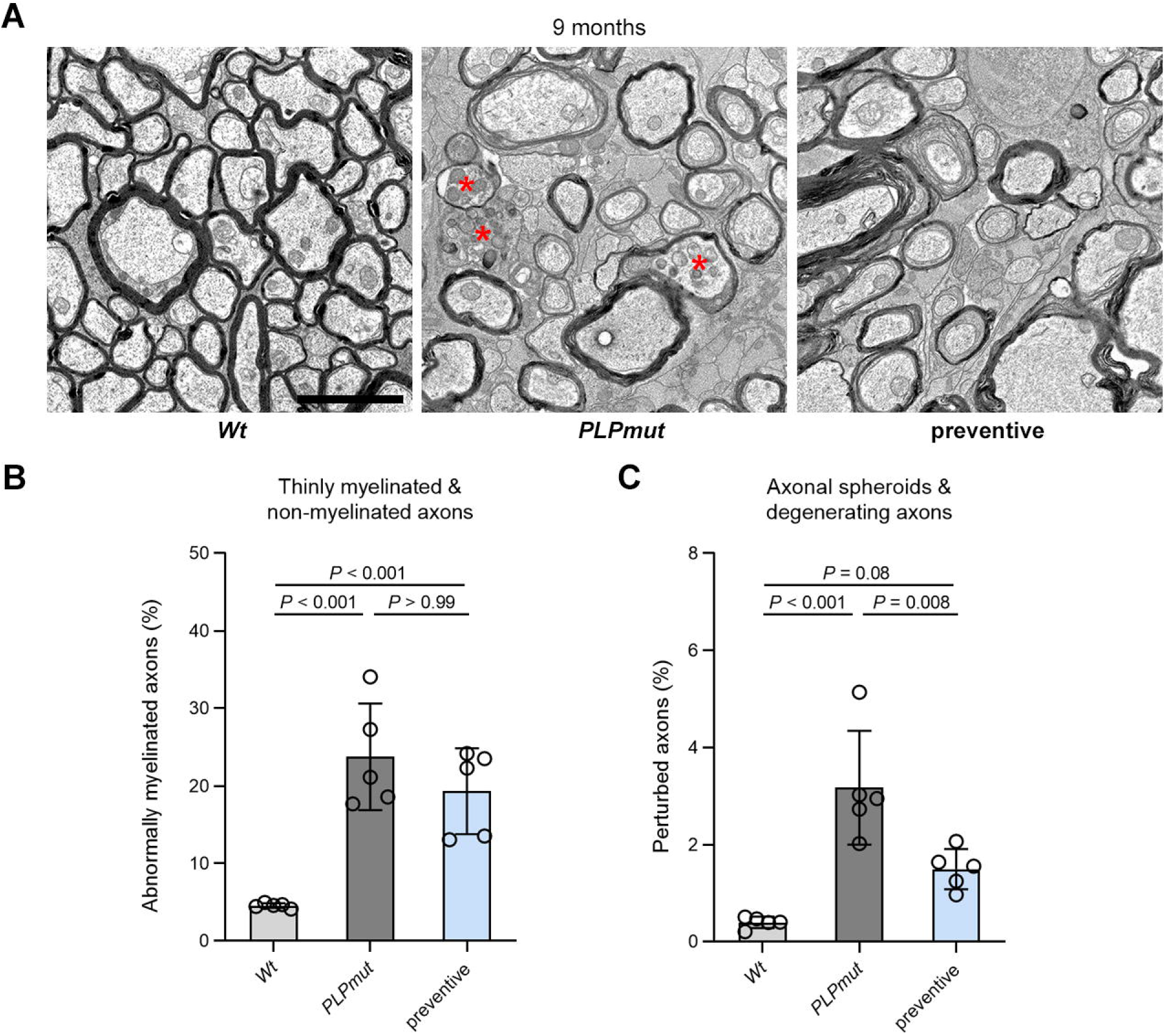
FTY720 treatment prevents T cell-driven axonal damage but not myelin pathology in *PLPmut* mice. (**A**) Representative electron micrographs of optic nerve cross-sections from *Wt*, *PLPmut*, and FTY720-treated *PLPmut* mice after the preventive treatment regimen indicated in Fig. 3 B. Asterisks indicate axons with focal accumulation of organelles. Scale bars, 2 μm. (**B**) Electron microscopy-based quantification of thinly myelinated (g-ratio ≥ 0.85) and non-myelinated axons or (**C**) axonal spheroids and degenerating axons in *Wt*, *PLPmut* and FTY720-treated *PLPmut* mice (*n* = 5 mice per group). Preventive FTY720 treatment attenuates axonal damage in *PLPmut* mice but has no effect on myelin alterations. B, C: one-way ANOVA with Tukey’s multiple comparisons test. Data are presented as the mean ± SD. All data represent at least three independent experiments.

To investigate putative direct neuroprotective effects of FTY720 treatment independent of adaptive immune reactions, we treated *PLPmut*/*Rag1*^−/−^ mice that lack mature adaptive immune cells and show mitigated but still detectable neurodegeneration^16^. Preventive treatment did not cause any additional amelioration of axon damage, neuron loss, and retinal thinning compared with untreated *PLPmut*/*Rag1*^−/−^ mice (Fig. S5).

### Axonal damage in PLPmut mice is driven by cytotoxic CD8^+^ T cells that specifically target mutant myelinating oligodendrocytes

Our previous data suggested a detrimental impact of the CD8^+^ T cell compartment in *PLPmut* mice^16,20,21^ and our unbiased analysis identified the strongest disease-related changes in CAT2, a cluster with the signature of tissue-resident memory and cytotoxic function. We therefore investigated the pathogenic role, putative cytotoxic effector mechanisms, and antigen/TCR dependency of CD8^+^ T cells by generating bone marrow chimeric *PLPmut* mice without confounding irradiation. *PLPmut*/*Rag1*^−/−^ mice lack mature adaptive immune cells and show diminished axonal damage^16^, enabling us to use them as recipients for comparing the impact of different adaptive immune cell populations/effector molecules. *PLPmut*/*Rag1*^−/−^ mice received bone marrow from various donor lines (Fig. 6 A), which leads to efficient and persistent restoration of adaptive immune cells within host tissues including the CNS^17,19,25,28^. Transplantation of *Wt* bone marrow into *Rag1* deficient *PLPmut* mice re-established T cell recruitment and the formation of SMI32^+^ axonal spheroids and thinning of the inner retina as assessed by OCT (Fig. 6 B-F). In contrast, *PLPmut*/*Rag1*^−/−^ mice reconstituted with *Cd8*^−/−^ bone marrow retained diminished axonal damage despite the presence of CD4^+^ T cells in the white matter. Similarly, reconstitution with *Gzmb*^−/−^ or *OT-I* (TCR specificity against ectopic ovalbumin) bone marrow did not result in more axonal spheroid formation and retinal thinning than in *PLPmut*/*Rag1*^−/−^ mice despite the restored recruitment of (genetically modified) T cells. In combination, these observations demonstrate a detrimental role of CD8^+^ T cells which damage axons in a granzyme B- and cognate TCR-dependent manner in *PLPmut* mice. Moreover, the normal recruitment but lack of impact of CD8^+^ T cells in BMC *OT-I* mice indicates white matter antigen specific activation.

**Figure 6.**
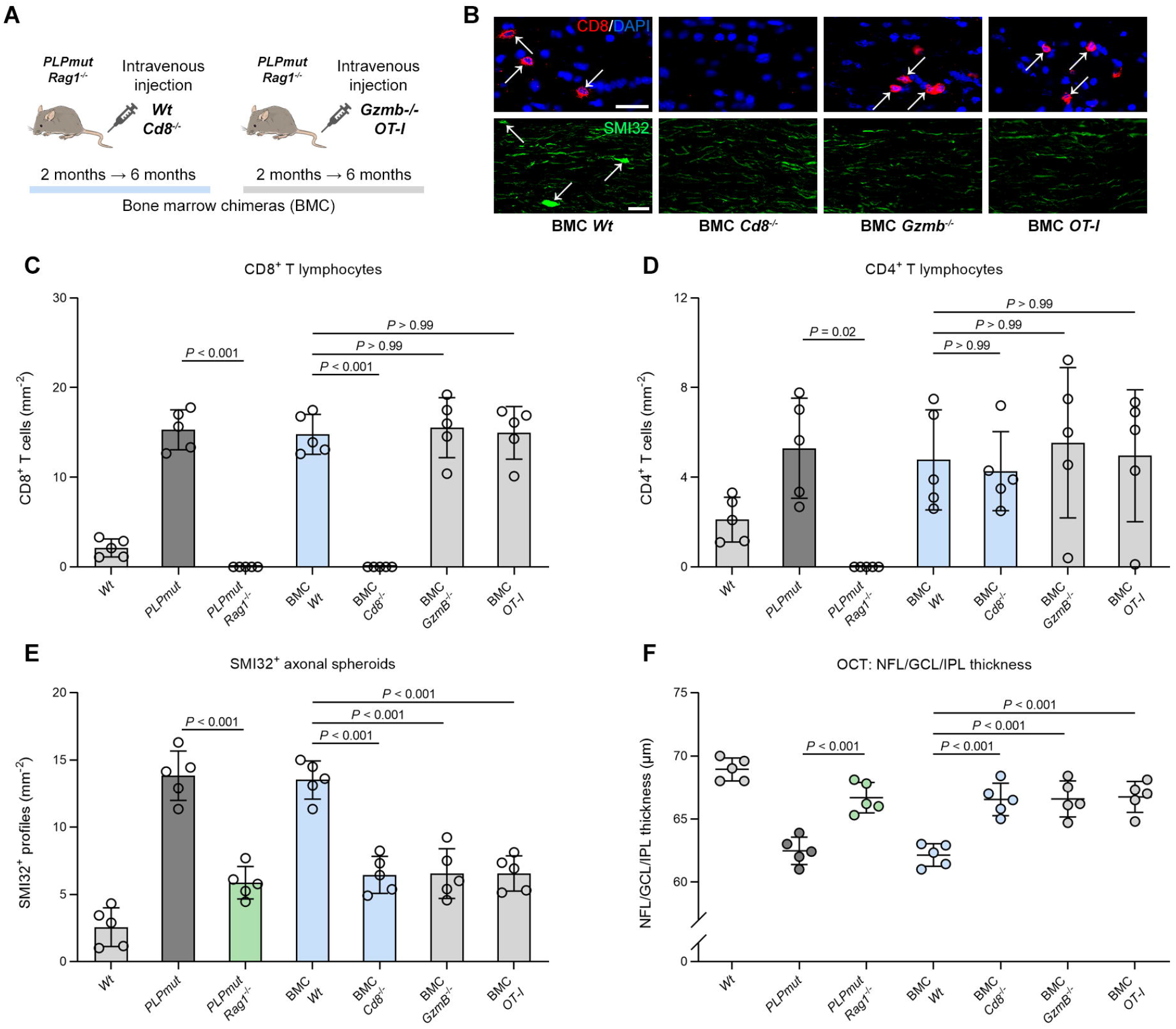
Myelinated axons in *PLPmut* mice are damaged by CD8^+^ T cells in a granzyme B and cognate TCR-dependent manner. (**A**) Schematic experimental design for generation of distinct *PLPmut*/*Rag1^−/−^* bone marrow chimeras (BMC) transplanted with bone marrow from *Wt*, *Cd8*^−/−^, *Gzmb*^−/−^ or *OT-I* mice. (**B**) Immunofluorescence detection of CD8^+^ T lymphocytes (top; arrows) as well as SMI32^+^ axonal spheroids (bottom; arrows) in the optic nerves of BMC. Scale bars, 20 μm. (**C**) Quantification of CD8^+^ T cells, (**D**) CD4^+^ T cells, and (**E**) SMI32^+^ axonal spheroids in the optic nerves from *Wt*, *PLPmut* and *Rag1*-deficient *PLPmut* mice as well as BMC transplanted with bone marrow from *Wt*, *Cd8*^−/−^, *Gzmb*^−/−^ or *OT-I* mice (*n* = 5 mice per group). (**F**) OCT analysis of the innermost retinal composite layer (NFL/GCL/IPL) in peripapillary circle scans in *Wt*, *PLPmut* and *Rag1*-deficient *PLPmut* mice as well as BMC (*n* = 5 mice per group). Axonal spheroid formation and retinal thinning is reestablished in *PLPmut*/*Rag1*^−/−^ BMC *Wt* mice but not in *PLPmut*/*Rag1*^−/−^ BMC *Cd8*^−/−^, BMC *Gzmb*^−/−^ and BMC *OT-I* mice. C-F: one-way ANOVA with Tukey’s multiple comparisons test. Data are presented as the mean ± SD. All data represent at least three independent experiments.

Since neuroinflammation in *PLPmut* mice results from gene defects specifically affecting oligodendrocytes, we wondered if CD8+ T cells target axon segments enwrapped by mutant myelin-producing cells. Due to random X chromosome inactivation, heterozygous female mice contain both *Wt* and *PLPmut* oligodendrocytes in the same white matter tracts. We detected a similar accumulation of CD8^+^ T cells but decreased Sn expression on microglia in optic nerves of heterozygous compared with homozygous females (Fig. 7 A, B, and C). Moreover, SMI32^+^ axonal spheroids were approximately half as frequent in mosaic mice (Fig. 7 D). *PLP1* mutations result in ultrastructural myelin compaction defects, which can be appreciated by electron microscopy^16^. We observed that spheroids and degenerating profiles in heterozygous females were almost exclusively surrounded by myelin segments displaying compaction defects, suggesting their formation by mutant oligodendrocytes (Fig. 7 E; 85.4% mutant myelin, 4.2% wt myelin, 10.4% no myelin; *n* = 100 damaged axons in 5 mice). Since loss of RGCs and retinal thinning were also halved by oligodendrocyte mosaicism (Fig. 7 F and G), we conclude that cytotoxic CD8^+^ T cells specifically target mutant myelinating oligodendrocytes which indirectly causes axonal damage and neurodegeneration. This matched with increased expression of the cognate antigen-presenting complex for CD8^+^ T cells - major histocompatibility complex class I (MHC-I) - on oligodendrocytes and microglia, especially in homozygous *PLPmut* mice (Fig. S6, Fig 7 H and I). To further investigate direct interactions between CD8^+^ T cells and *Wt* and *PLPmut* oligodendrocytes, we analyzed citrullination of MBP, which is typically enhanced in perturbed and destabilized myelin and increases its susceptibility to inflammation^23^. Indeed, citrullinated MBP was strongly enriched in optic nerves of *PLPmut* mice and was less homogenously distributed in heterozygous *Wt*/*PLPmut* mice (Fig. 7 J). Using triple immunofluorescence, we found that an association of CD8^+^ T cells with SMI32^+^ damaged axons was almost exclusively detectable at segments ensheathed by *PLPmut* (citrullinated MBP^+^) myelin (Fig. 7 K and L).

**Figure 7.**
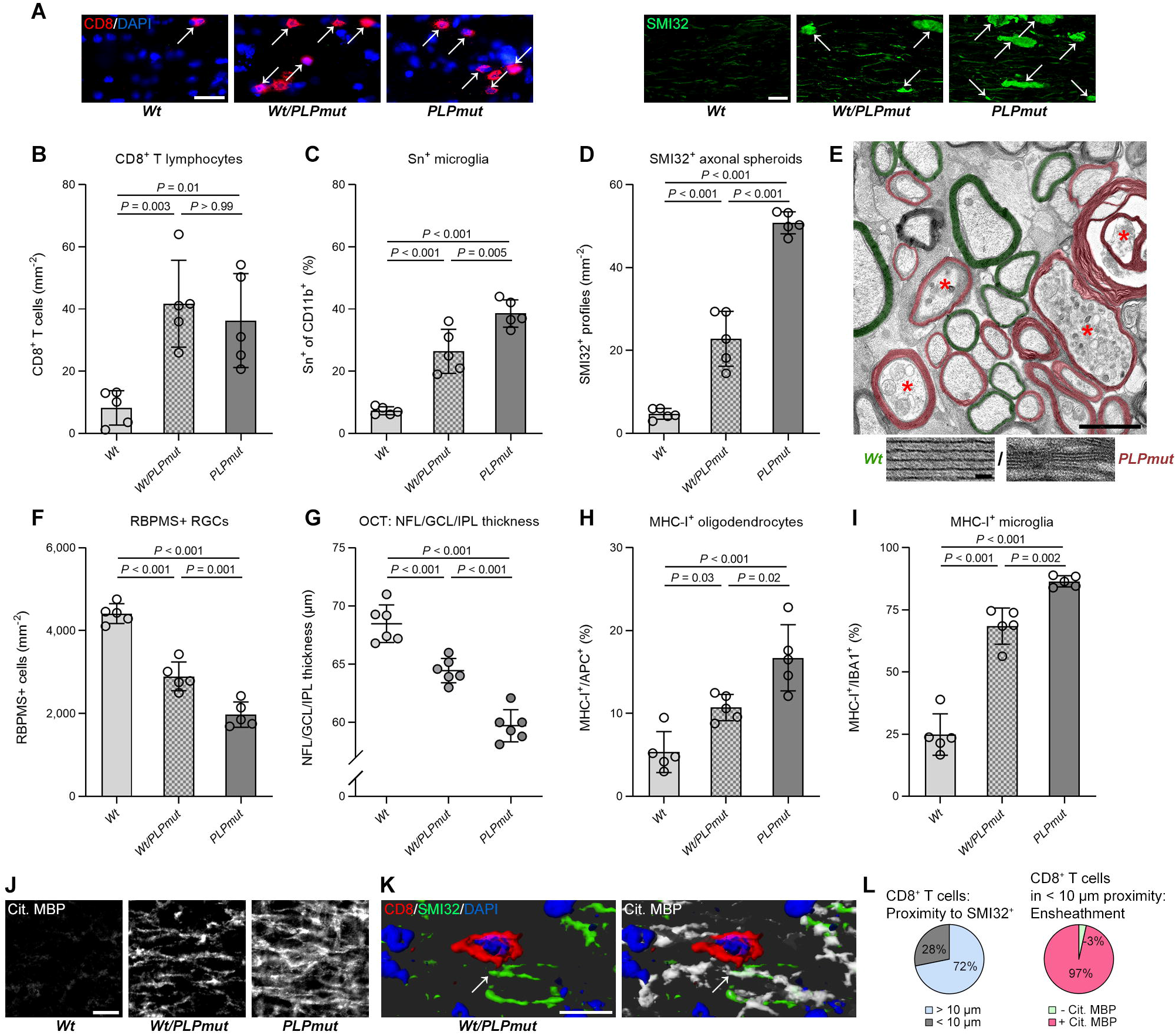
CD8^+^ T cells selectively damage axon segments myelinated by *PLP1* mutant oligodendrocytes. (**A**) Immunofluorescence detection of CD8^+^ T lymphocytes (left; arrows) as well as SMI32^+^ axonal spheroids (right; arrows) in the optic nerves from 18-month-old female *Wt*, *Wt*/*PLPmut* (heterozygous), and *PLPmut* (homozygous) mice. Scale bars, 20 μm. (**B**) Quantification of CD8^+^ T cells, (**C**) activated Sn^+^ microglia (% of CD11b^+^), and (**D**) SMI32^+^ axonal spheroids in the optic nerves of 18-month-old female *Wt*, *Wt*/*PLPmut* (heterozygous), and *PLPmut* (homozygous) mice (*n* = 5 mice per group). There is a similar density of CD8^+^ T cells, but less microglial activation and axonal damage in white matter of mosaic mice. (**E**) Representative electron micrograph of an optic nerve cross-section from a *Wt*/*PLPmut* (heterozygous) female. Asterisks indicate axons with focal accumulation of organelles or degenerating axons. Pseudocolor indicates normal (green) or perturbed, non-compacted (magenta) myelin segments formed by *Wt* or *PLPmut* oligodendrocytes, respectively. Scale bar, 2 μm. Representative high-resolution images (bottom) of *Wt* and *PLPmut* myelin demonstrate collapsed or vanishing intraperiod lines within mutant myelin. Scale bar, 20 nm. (**F**) Quantification of RGCs and (**G**) OCT analysis of the innermost retinal composite layer (NFL/GCL/IPL) in peripapillary circle scans in female *Wt*, *Wt*/*PLPmut* (heterozygous), and *PLPmut* (homozygous) mice (*n* = 5 mice per group). Neuron loss and retinal thinning is lower in mosaic mice. (**H**) Quantification of MHC-I^+^ oligodendrocytes (% of APC^+^) and (**I**) MHC-I^+^ microglia (% of Iba1^+^), in the optic nerves of female *Wt*, *Wt*/*PLPmut* (heterozygous), and *PLPmut* (homozygous) mice (*n* = 5 mice per group). MHC-I expression on oligodendrocytes and microglia is lower in mosaic mice. (**J**) Immunofluorescence detection of citrullinated MBP in the optic nerves from 18-month-old female *Wt*, *Wt*/*PLPmut* (heterozygous), and *PLPmut* (homozygous) mice. Scale bar, 20 μm. (**K**) Immunofluorescence detection and IMARIS-based Z-Stack reconstruction of CD8, SMI32, and citrullinated MBP in the optic nerves from 18-month-old *Wt*/*PLPmut* (heterozygous) mice. Scale bar, 10 μm. (**L**) Quantifications show that the association (< 10 μm proximity) of CD8^+^ T cells to SMI32^+^ damaged axons (arrow in panel K) occurs almost exclusively when they are ensheathed by mutant (cit. MBP^+^) myelin (*n* = 103 CD8^+^ T cells from 5 mosaic female mice. B-D and F-I: one-way ANOVA with Tukey’s multiple comparisons test. Data are presented as the mean ± SD. All data represent at least three independent experiments.

## Discussion

Recent studies have renewed the appreciation of oligodendrocyte-lineage cells as immunocompetent glial cells that can initiate or perpetuate neuroinflammation ^5,24,29–32^. This has relevance for inherited and acquired diseases of myelin and supports the hypothesis of CNS-intrinsic perturbation as one possible mechanism (among multiple other factors) contributing to chronic adaptive immune reactions with a detrimental impact on white matter integrity^33–35^. We have previously shown that defects in myelin genes result in secondary neuroinflammation that contributes to axonal damage and functional impairment^16,17^. *PLP1* mutant oligodendrocytes carry point mutations that have been identified in patients with multiple sclerosis and seem to acquire disease-associated characteristics at least partially shared (e.g., increased MHC-I expression) with other diseases and normal aging^11,32,36^. Moreover, normal aging is also associated with an accumulation of CD8^+^ T cells in the white matter that appear to target myelinated axons and contribute to neurodegeneration, cognitive, and motor decline^25^. These observations reflect the vulnerability of myelinated axons to immune-mediated perturbation upon various primary defects and underscore an active role of myelinating glia in these conditions. Moreover, targeting pro-inflammatory microglia in *PLPmut* mice attenuates T cell-driven axon damage^20^, supporting the hypothesis that disturbed glial interactions play a major part in the initiation of neuroinflammation.

Previous work has indicated that CD8^+^ T cells populating the CNS parenchyma resemble tissue-resident memory (TRM) cells which do not recirculate but respond to peripheral stimuli^37–39^. Our characterization of these cells in aging^25^ and myelin mutant mice (this study) is in line with this and revealed that different subpopulations of these cells show signatures related to maintenance and long-term residency within the CNS. While our exploratory scRNA-seq approach required pooled analysis of rare CD8^+^ T cells from brains of multiple mice (limiting insights into variation), we validate our major conclusions using independent techniques and confirm the phenotype, heterogeneity, and disease-related changes of these cells. A hallmark of TRM cells is the downregulation of receptors involved in tissue egress and recirculation once they have infiltrated their respective niches^40,41^. Consequently, TRM cells become resistant to S1PR modulation and systemic depletion of T cells has little effect on frequencies and impact within the infected or inflamed tissue. Our distinct treatment regimens with fingolimod confirm that early (preventive) treatment causes lymphopenia and attenuates CNS recruitment of T cells, whereas late (therapeutic) treatment still depletes circulating T cells but has no effect on those associated with the CNS parenchyma. The relative stability of CD8^+^ T cell densities but accelerating accumulation of axonal damage at advanced disease stages might indicate either earlier compensatory/resilience mechanisms or a pronounced detrimental impact of long-term resident T lymphocytes with limited turnover. Recent observations support a local contribution of CD8^+^ TRM cells to chronic CNS autoimmunity^42,43^. In multiple sclerosis, active lesions are dominated by CD8^+^ T cells that infiltrate the parenchyma, partially acquire features of tissue-resident memory cells, and persist in the CNS^44^. Activated, clonally expanded CD8^+^ TRM cells are also present in the CSF from MS-discordant monozygotic twins with subclinical neuroinflammation (prodromal multiple sclerosis)^45^. Interestingly, these cells at least partially express CD103 and accumulate in normal-appearing white matter of MS patients, where they might sustain diffuse chronic inflammation and axonal damage especially in progressive forms of the disease^46–48^. Here we observed using multiple independent methods that a subset of CD8^+^ CAT with the highest expression of CD103 (CAT2) shows the most prominent disease-related changes in *PLPmut* mice, indicating that the myelin mutant model is suitable to study neural-immune interactions with relevance for chronic neuroinflammatory diseases.

The same population also showed the highest expression of *Gzmb* among parenchymal CD8^+^ T cells, indicating that like in aging and *PLPtg* mice^19,25^, axonal damage might depend on the cytolytic effector protease. By bone marrow transfer into *Rag1*-deficient *PLPmut* mice, we demonstrate that CNS-infiltrating CD8^+^ T cells require *Gzmb* expression and cognate TCR specificity (BMC *OT-I*) to drive axonal damage. In contrast, reconstitution of CD4^+^ T cells and B cells in the absence of CD8^+^ T cells (BMC *Cd8*^−/−^) does not result in more axonal damage than in genuine *PLPmut*/*Rag1*^−/−^ mice (displaying attenuated axonal damage). These reconstituted CD8^−^ lymphocytes are unable to damage myelinated axons despite otherwise similar characteristics (CNS infiltration, putative effector molecule expression) as in genuine *PLPmut* mice. Thus, axonal damage in *PLPmut* mice appears to be driven by antigen specific, autoreactive CD8^+^ T cells that use contact-dependent (cytotoxic granules) effector mechanisms. The similarity of these processes between distinct myelin mutants and normal white matter aging is a novel insight and indicates a conserved feature possibly shared with many other conditions. While the exact antigen(s) recognized by these cells remains to be determined in future studies, several observations lead us to speculate that they might target perturbed myelin. First, CD8^+^ T cells are predominantly accumulating in white matter tracts and associate with juxtaparanodal domains of mutant fibers^16^, similar as previously observed in *PLPtg* mice^18^. Second, axons showing focal signs of damage and degeneration are almost exclusively enwrapped by perturbed mutant myelin in heterozygous (mosaic) females. Third, myelin mutant oligodendrocytes express increased levels of MHC-I compared with healthy (Wt) myelinating glia, indicating that they might show increased (auto)antigen presentation to CD8^+^ T cells. It is conceivable that myelin gene defects (and aging) result in molecular changes and cell stress pathways in oligodendrocyte subsets that initiate their communication with immune cells and make them susceptible to immune-mediated damage, similar as recently shown for selective neuron populations in AD^49^. Among those glial changes is enhanced citrullination of MBP, as occurs in destabilized myelin after toxin- or myelin disease-related perturbation^23,50^. We found that CD8^+^ T cells in proximity to SMI32^+^ axons were almost exclusively associated with segments showing enhanced MBP citrullination, again indicating specificity for mutant myelin-related components. Such changes in myelin properties might lead to its recognition as neoantigen when presented to T cells.

Considering that a cytotoxic T cell attack on perturbed myelinating oligodendrocytes drives axonal damage, it remains to be clarified how this impairment is mediated before obvious demyelination. Collateral damage, changes in trophic/metabolic support of axons, or other detrimental reactions of myelinating oligodendrocytes in response to T cell-derived lytic granules might be among the responsible mechanisms^51–53^. In aged mice, scRNA-seq has revealed some of the transcriptional changes in oligodendrocytes upon being targeted by CD8^+^ T cells^32^. Both the frequencies of *Serpina3n*^+^ (an inhibitor of proteases including granzyme B) and interferon-responsive oligodendrocytes in the white matter were decreased upon *Rag1* deficiency, indicating that oligodendrocytes show distinct responses to neuroinflammation. We speculate that this might be related to the impact of *Gzmb*-vs *Ifng*-expressing CD8^+^ T cell populations and have differential effects on axon-glia interactions and axon degeneration. Our previous characterization of CD8^+^ CAT from aged mice revealed the strong increase of a specific population expressing checkpoint molecules that is not responding in *PLPmut* mice and a downregulation of *Ifng*^25^. Combined with the observation that axonal damage in aged and myelin mutant mice depends on granzyme B, we propose that the population of oligodendrocytes upregulating *Serpina3n* to counteract T cell cytotoxicity, demyelination, and cell death is more detrimental to axonal integrity. Indeed, axons that remain myelinated under inflammatory conditions have recently been proposed to be at higher risk for degeneration^54^. The fact that FTY720 treatment attenuated T cell recruitment and axonal damage but did not significantly affect myelin integrity in *PLPmut* mice also argues against the possibility of axon degeneration being a consequence of demyelination. Since a direct T cell attack on axons is also difficult to explain when considering the reduced damage in mosaic females, further studies should explore the detrimental T cell-driven reactions of *PLPmut* oligodendrocytes.

Our observations are of translational relevance for neurological conditions associated with myelin defects and chronic neuroinflammation and show that early onset of therapy might be critical for the efficacy of S1PR-modulation to prevent recruitment and CNS colonization of T cells. When these cells become resident, they downregulate receptors for tissue egress and appear to stay within the CNS for extended time periods without relying on high turnover from the circulation. Moreover, cessation of circulating lymphocyte ablation in *PLPmut* mice leads to a slow restoration of T cells within the CNS with sustained benefits on neural integrity, reflecting the low-grade accumulation of inflammatory damage typical of chronic disease. The lack of any beneficial effect upon therapeutic treatment regarding the progression of neurodegeneration argues against a direct protective effect of fingolimod in *PLPmut* mice. Indeed, FTY720 treatment of *PLPmut*/*Rag1*^−/−^ mice (lacking adaptive immune cells and showing much milder - but still detectable - pathology than genuine *PLPmut* mice) did not cause any additional amelioration of axon damage and neuron loss. This is supported by our previous treatment approaches in *Rag1*-deficient models of rare lysosomal storage diseases accompanied by T cell-driven axon degeneration, which also did not reveal major effects independent of immune cells^55^.

The resilience of CNS-associated T cells and compartmentalized inflammatory response might explain the limited efficacy of a previous clinical trial using fingolimod in primary progressive multiple sclerosis^56^. Furthermore, they implicate that neuroaxonal degeneration in such chronic conditions might still be related to low-grade inflammation, going along with the poor efficacy of S1PR-modulating therapies. Interestingly, we previously observed that late onset therapeutic treatment with teriflunomide, another established drug for MS, showed beneficial effects and halted ongoing axon degeneration in *PLPmut* mice^21^. Teriflunomide is an inhibitor of dihydroorotate dehydrogenase and modulates mitochondrial respiration, T cell activation and migration, particularly within the CD8^+^ compartment^57^. Moreover, it induces a tolerogenic bias in immune cells of MS patients^58^. In the myelin mutants, therapeutic treatment with teriflunomide resulted in an increased frequency of CD8^+^ T cells expressing high levels of CD122 and PD-1 in the white matter^21^. Such cells have been shown to restrict inflammation and autoimmunity by suppressing effector T cells, and regulatory populations within the CD8 lineage have been described in mice and humans^59–64^. Combining our previous and present observations, we speculate that CD8^+^ CAT5 might represent such an anti-inflammatory population within the CNS. This cluster is enriched with marker genes of regulatory function (Fig. 2 A) and shows low expression of *Txnip* (enriched in CAT2; Table S1), a sensor of oxidative phosphorylation and glycolysis^65^, which might explain its resilience to inhibition of mitochondrial metabolism. Therapeutic strategies to modulate the activation or change the composition of CD8^+^ CNS-associated T cells towards tolerance might be preferred over more general immunosuppression. Moreover, our data indicate that treatment time point/disease stage and activity must be carefully considered when selecting specific immunomodulatory treatment approaches for chronic neuroinflammation. Nevertheless, we here emphasize that targeting neuroinflammation might be a feasible approach for chronic progressive degenerative diseases associated with myelin defects and aging.

## Acknowledgements

We thank H. Blazyca, S. Loserth and B. Meyer for technical assistance and J. Schreiber, A. Weidner, and T. Bimmerlein for their attentive care of mice. This work was supported by Novartis Pharma GmbH (Germany), the German Research Foundation (grant no. MA1053/6-2 to R.M. and grant no. GR5240/1-1 to J.G.), the Roman, Marga und Mareille Sobek Foundation (to R.M.), the Charitable Hertie Foundation (grant no. P1150084 to J.G.) and the Interdisciplinary Centre for Clinical Research of the University of Würzburg (grant no. A-302 to R.M. and grant no. Z-6 to P.A.).

## Author contributions

J.G. and R.M. planned and oversaw all aspects of the study. T.A., D.S., and J.G. performed and analyzed most of the experiments. K.K. and W.K. performed flow cytometry and analyzed the scRNA-seq experiments. P.A. and A.E.S. performed cell sorting and scRNA-seq. J.G. wrote the manuscript with input from all authors.

## Methods

### Resource availability

#### Lead contact

Further information and requests for resources and reagents should be directed to and will be fulfilled by the lead contact, Janos Groh.

#### Materials availability

This study did not generate new unique reagents.

#### Data and code availability

- The single cell RNA sequencing data in this publication have been deposited in the Gene Expression Omnibus (GEO): GSE138891. Other data that support the findings of this study will be shared by the lead contact upon request.
- This paper does not report original code.
- Any additional information required to reanalyze the data reported in this paper is available from the lead contact upon request.

### Experimental model and subject details

Mice were kept at the animal facility of the Centre for Experimental Molecular Medicine, University of Würzburg, under barrier conditions and at a constant cycle of 14 h in the light (<300 lux) and 10 h in the dark. Colonies were maintained at 20-24 °C and 40-60% humidity, with free access to food and water. All animal experiments were approved by the Government of Lower Franconia, Germany. All mice including Wt (*Wt*), *PLPmut* (B6.Cg-Tg(PLP1)1Rm*-Plp1tm1Kan*/J)^16^ - genuine or crossbred with *Rag1*^−/−^ (B6.129S7-*Rag1tm1Mom*/J)^66^, *Cd8*^−/−^ (B6.129S2-*Cd8atm1Mak*/J), *Gzmb*^−/−^ (B6.129S2-*Gzmbtm1Ley*/J)^67^, *OT-I* (C57BL/6-Tg(TcraTcrb)1100Mjb/J)^68^ mice were on a uniform C57BL/6J genetic background; they were bred, regularly backcrossed and aged in-house. Since we did not detect obvious differences between male and female mice in the analyses presented in the current study, mice of either sex were used for most of the experiments (except for analyzing heterozygous vs homozygous females). Genotypes were determined by conventional PCR using isolated DNA from ear punch biopsies.

### Method details

#### Flow cytometry and cell sorting

Mice were euthanized with CO_2_ (according to the guidelines by the State Office of Health and Social Affairs Berlin) and blood was thoroughly removed by transcardial perfusion with PBS containing heparin. Brains including optic nerves, leptomeninges and choroid plexus were dissected, collected in ice-cold PBS and cut into small pieces. Tissue was digested in 1 ml of Accutase (Merck Millipore) per brain at 37 °C for 30 min and triturated through 100-μm cell strainers, which were rinsed with 10% FCS in PBS. Cells were purified by a linear 40% Percoll (GE Healthcare) centrifugation step at 650 g without brakes for 25 min and the myelin top layer and supernatant were discarded. Mononuclear cells were resuspended in fluorescence-activated cell sorting buffer (1% BSA and 0.1% sodium azide in PBS) and isolated cells were counted for each brain. For scRNA-seq, cells from the brains of 5 adult (12-month-old) *Wt* and 4 *PLPmut* mice were pooled into 2 separate samples. Pooled samples were processed in parallel to avoid batch effects. Viable cells were identified by LIVE/DEAD stain (catalog no. L34965; Thermo Fisher Scientific), Fc receptors were blocked for 15 min with rat anti-CD16/32 (1:100, catalog no. 553141; BD Biosciences) and cells were washed and labeled with the following antibodies for 30 min at 4 °C: rat anti-CD8 APC (1:100, catalog no. 553035; BD Biosciences); rat anti-CD45 PE (1:100, catalog no. 103105; BioLegend). Cells were washed twice, single viable cells were gated and CD45^high^CD8^+^ cells were collected using a FACSAria III and corresponding software (FACSDiva, v.6; BD Biosciences). Calculation of the number of CD45^high^CD8^+^ T cells per brain was performed by extrapolating their frequency to the counted total number of isolated cells. For further experiments viable CD45^high^CD8^+^ cells were labeled with rat anti-CD45 APC (1:100, catalog no. 103111; BioLegend), rat anti-CD8 PerCP/Cyanine5.5 (1:100, catalog no. 100733; BioLegend), rat anti-CXCR6 PE/Cyanine7 (1:100, catalog no. 151118; BioLegend), rat anti-CXCR4 PE (1:100, catalog no. 146505; BioLegend), armenian hamster anti-CD103 BV605 (1:100, catalog no. 121433; BioLegend), and rat anti-Ly6A/E FITC (1:100, catalog no. 108105; BioLegend). Cells were washed twice; single viable cells were gated and CD45^high^CD8^+^ cells were analyzed using a FACSLyric (BD Biosciences) and FlowJo (version 10; LLC). Contribution of the different subsets in absolute numbers was calculated by extrapolating their frequencies to the number of CD45^high^CD8^+^ T cells per brain. Circulating leukocytes were quantified in peripheral blood samples. Before transcardial perfusion, blood was collected from the right atrium of the heart using a heparinized capillary and coagulation was prevented by adding PBS containing heparin. Erythrocytes were lysed and the remaining cells were washed and analyzed by flow cytometry. Total leukocytes were gated based on forward and side scatter, myeloid cells were stained using PE-conjugated antibodies against CD11b (1:100, catalog no. 557397; BD Biosciences), and T lymphocytes were stained using antibodies against CD4 and CD8 (1:100, catalog nos. 553049 and 553032; BD Biosciences). At least 1◻×◻10^5^ leukocytes per mouse were analyzed and their amount per microliter of blood was calculated.

#### Single-cell RNA sequencing (scRNA-seq) and data processing

Around 15,000 CD45^high^CD8^+^ single cells were sorted per sample using a FACSAria III (BD Biosciences) before being encapsulated into droplets with the Chromium Controller (10x Genomics) and processed according to the manufacturer’s specifications. Briefly, every transcript captured in all the cells encapsulated with a bead was uniquely barcoded using a combination of a 16-base pair (bp) 10x barcode and a 10-bp unique molecular identifier (UMI). Complementary DNA libraries ready for sequencing on Illumina platforms were generated using the Chromium Single Cell 3′ Library & Gel Bead Kit v2 (10x Genomics) according to the detailed protocol provided by the manufacturer. Libraries were quantified by Qubit 3.0 Fluorometer (Thermo Fisher Scientific) and quality was checked using a 2100 Bioanalyzer with High Sensitivity DNA kit (Agilent Technologies). Libraries were pooled and sequenced with a NovaSeq 6000 platform (S1 Cartridge; Illumina) in paired-end mode to reach a mean of 75,412 reads per single cell. A total of 5,017 and 6,467 cells were captured and a median gene number per cell of 1,325 and 1,283 could be retrieved for adult *Wt* and *PLPmut* cells, respectively. Data were demultiplexed using the CellRanger software v.2.0.2 based on 8 bp 10x sample indexes; paired-end FASTQ files were generated. The cell barcodes and transcript unique molecular identifiers were processed as described previously^69^. The reads were aligned to the University of California, Santa Cruz mouse mm10 reference genome using STAR aligner^70^ v.2.5.1b. The alignment results were used to quantify the expression level of mouse genes and generate the gene-barcode matrix. The cellranger aggr command of CellRanger was used to aggregate different libraries. Subsequent data analysis was performed using the R package Seurat^71^ v.2.4 and 4.0. Doublets and potentially dead cells were removed based on the percentage of mitochondrial genes (cutoff set at 5%) and the number of genes (cells with >800 and <2,200 genes were used) expressed in each cell as quality control markers. The gene expression of the remaining cells (4,338 and 5,110 cells from *Wt* and *PLPmut* mice, respectively) was log-normalized. Highly variable genes were detected with Seurat and the top 1,000 of these genes were used as the basis for downstream clustering analysis after regressing out mitochondrial expression per cell. Principle component analysis was used for dimensionality reduction and the number of significant principal components was calculated using the built-in JackStraw function. Cells were clustered based on the identified principal components (16) with a resolution of 0.6; uniform manifold approximation and projection was used for data visualization in two dimensions. A minimal contamination of myeloid cells was removed based on marker gene expression. Contribution of the samples to each cluster in absolute numbers was calculated by extrapolating their frequencies to the number of CD45^high^CD8^+^ T cells per brain. Differentially expressed genes were identified with min.pct = 0.25 and a cutoff of p_val_adj > 0.05. Complete lists of differentially expressed genes are included in Table S1. Marker gene scores for feature expression programs were calculated using the AddModuleScore function in Seurat.

#### Immunomodulatory treatment

Fingolimod (FTY720, provided by Novartis, Basel, Switzerland) was dissolved in autoclaved drinking water at 3 μg/mL and provided *ad libitum*. With an approximate consumption of 5 ml/day and 30 g body weight, this corresponds to a dose of 0.5 mg/kg body weight/day. This concentration is based on previous animal experiments^72,73^ and approximately corresponds to doses used for human multiple sclerosis patients, when a dose conversion scaling is applied^74^. Non-treated controls received autoclaved drinking water and the water with or without FTY720 was changed weekly. Mice were treated for 75 or 150 days and monitored daily regarding defined burden criteria and phenotypic abnormalities. No obvious side effects or significant changes in body weight were detected upon the treatment.

#### Histochemistry and immunofluorescence

Mice were euthanized with CO_2_ (according to the guidelines by the State Office of Health and Social Affairs Berlin), blood was removed by transcardial perfusion with PBS containing heparin and tissue was fixed by perfusion with 2% paraformaldehyde (PFA) in PBS. Tissue was collected, postfixed, dehydrated, and processed as described previously^16^. Blood was collected before transcardial perfusion from the right atrium using a heparinized capillary and smears were air-dried overnight. Immunohistochemistry was performed on 10-μm-thick longitudinal optic nerve or spleen cryo-sections and blood smears after postfixation in 4% PFA in PBS or ice-cold acetone for 10 min. Sections were blocked using 5% BSA in PBS and incubated overnight at 4 °C with 1 or an appropriate combination of up to 3 of the following antibodies: rat anti-CD8 (1:500, catalog no. MCA609G; Bio-Rad Laboratories), rat anti-CD4 (1:1,000, catalog no. MCA1767; Bio-Rad Laboratories), rat anti-CD8 biotinylated (1:500, catalog no. 553028; BD Biosciences), rabbit anti-LAG3 (1:100, catalog no. ab209238; Abcam), rat anti-CXCR4 PE (1:100, catalog no. 146505; BioLegend), armenian hamster anti-CD103 BV605 (1:100, catalog no. 121433; BioLegend), rat anti-Ly6A/E FITC (1:100, catalog no. 108105; BioLegend), rat anti-CD11b (1:100, catalog no. MCA74G; Bio-Rad Laboratories), rabbit anti-CD11b (1:100, catalog no. NB110-89474; Novus Biologicals), rat anti-Sn (1:300, catalog no. MCA947G; Bio-Rad Laboratories), hamster anti-CD11c (1:100, catalog no. MA11C5; Thermo Fisher Scientific), rabbit anti-P2RY12 (1:300, catalog no. 55043A, AnaSpec), mouse anti-SMI32 (1:1,000, catalog no. 801701; BioLegend), rat anti-MHC-I (1:100, catalog no. T-2105; Dianova), goat anti-Iba1 (1:300, catalog no. NB100-1028; Novus Biologicals), mouse anti-APC (1:300, catalog no. ab16794; Abcam), rabbit anti-citrullinated MBP (1:300, catalog no. 26742; Cayman Chemical); For indirect detection, immunoreactive profiles were visualized using fluorescently labeled (1:300; Dianova) secondary antibodies, streptavidin (1:300; Thermo Fisher Scientific) or biotinylated secondary antibodies (1:100; Vector Laboratories) and streptavidin-biotin-peroxidase (Vector Laboratories) complex using diaminobenzidine HCl and H_2_O_2_; nuclei were stained with 4,6-diamidino-2-phenylindole (DAPI) (Sigma-Aldrich). Light and fluorescence microscopy images were acquired using an Axio Imager M2 microscope (ZEISS) with ApoTome.2 structured illumination equipment, attached Axiocam cameras and corresponding software (ZEN v.2.3 blue edition) or a FluoView FV1000 confocal microscope (Olympus) with corresponding software (v.2.0). Images were minimally processed (rotation, cropping, addition of symbols) to generate figures using Photoshop CS6 and Illustrator CS6 (Adobe). For quantification, immunoreactive profiles were counted in at least three nonadjacent sections for each animal and related to the area of these sections using the cell counter plugin in Fiji/ImageJ v.1.51 (National Institutes of Health). To quantify RGCs, perfusion-fixed eyes were enucleated, and specific markers of the inner retinal cell types were labeled in free-floating retina preparations. Fixed retinae were frozen in PBS containing 2% Triton X-100, thawed, washed and blocked for 1 h using 5% BSA and 5% donkey serum in PBS containing 2% Triton X-100. Retinae were incubated overnight on a rocker at 4 °C with appropriate combinations of the following antibodies: guinea pig anti-RBPMS (1:300, catalog no. ABN1376; Merck Millipore); goat anti-Brn3a (1:100, catalog no. sc-31984; Santa Cruz Biotechnology); immune reactions were visualized using fluorescently labeled (1:500; Dianova) secondary antibodies, retinae were flat-mounted, and the total retinal area was measured. RGCs were quantified in three images of the middle retinal region per flat mount using the cell counter plugin in Fiji/ImageJ v.1.51 (National Institutes of Health). Images were taken at a fixed distance (~1 mm) and magnification from the optic nerve head in three different quadrants of the flat mounts.

#### Electron microscopy

The optic nerves of transcardially perfused mice were postfixed overnight in 4% PFA and 2% glutaraldehyde in cacodylate buffer. Nerves were osmicated and processed for light and electron microscopy; morphometric quantification of neuropathological alterations was performed as published previously^16^ using a LEO906 E electron microscope (ZEISS) and corresponding software iTEM v.5.1 (Soft Imaging System). At least 10 regions of interest (corresponding to an area of around 5% and up to 3,000 axons per individual optic nerve) were analyzed per optic nerve per mouse. The percentages of axonal profiles showing spheroid formation or undergoing degeneration were identified individually by their characteristic morphological features in electron micrographs and related to the number of all investigated axons per optic nerve per mouse. Genetically perturbed myelin in heterozygous females was identified by myelin compaction defects at high resolution. Images were processed (rotation, cropping, addition of symbols and pseudocolor) to generate figures using Photoshop CS6.

#### Spectral domain optical coherence tomography (OCT)

Mice were subjected to OCT imaging with a commercially available device (SPECTRALIS OCT; Heidelberg Engineering) and additional lenses as described previously^16,75^. Mice were measured at different ages for longitudinal analysis and the thickness of the innermost retinal composite layer comprising the nerve fiber layer (NFL), GCL and inner plexiform layer (IPL) were measured in high-resolution peripapillary circle scans (at least ten measurements per scan) by an investigator unaware of the genotype and treatment condition of the mice using HEYEX v.1.7.1.

#### Bone marrow transplantation

Bone marrow was transferred according to previously published protocols^25,76^. Briefly, bone marrow was isolated from the femur and tibia of donor mice and 1 × 10^7^ cells were injected intravenously into anaesthetized *PLPmut*/*Rag1*^−/−^ mice; this provides a niche for engraftment and long-term reconstitution of adaptive immune cells without confounding irradiation^28^. *PLPmut*/*Rag1*^−/−^ mice were reconstituted at 2 months of age and analyzed at 6 months of age. Successful chimerism was controlled by flow cytometry of splenocytes and immunohistochemistry on optic nerve sections. Engraftment of transplanted bone marrow led to a frequency of the respective T lymphocyte types in the *PLPmut*/*Rag1*^−/−^ hosts which was similar to *PLPmut*/*Rag1*^+/+^ mice.

## Quantification and statistical analysis

All quantifications and analyses were performed by blinded investigators who were unaware of the genotype and treatment group of the respective mice or tissue samples after concealment of genotypes with individual uniquely coded labels. Animals were randomly placed into experimental or control groups according to the genotyping results using a random generator (http://www.randomizer.org). For biometrical sample size estimation, the program G*Power v.3.1.3 was used^77^. Calculation of appropriate sample size groups was performed using an *a priori* power analysis by comparing the mean of 2 to 4 groups with a defined adequate power of 0.8 (1 - beta error) and an α error of 0.05. To determine the prespecified effect size d or f, previously published data were considered as comparable reference values^16,20,21^. This resulted in large prespecified effect sizes ranging from 1.20 to 3.56 for our primary outcome measures (densities of T cells, SMI32^+^ axonal spheroids, RBPMS^+^ RGCs, OCT of inner retinal thinning). The number of individual mice per group (number of biologically independent samples) for each experiment and the meaning of each data point are indicated in the respective figure legends. All data (except the scRNA-seq experiment) represent at least three independent experiments. For this, we quantified a specific cell type/structure in multiple different sections/samples of a respective tissue and averaged the measurements into one single data point. No animals were excluded from the analyses. In the scRNA-seq experiment, we had to pool the brains of 4–5 mice for each age/genotype group due to the low number of T cells in the CNS. Statistical analysis was performed using Prism 8 (GraphPad Software). The Shapiro-Wilk test was used to check for the normal distribution of data and the F test was used to check the equality of variances to ensure that all data met the assumptions of the statistical tests used. Comparisons of two groups were performed with an unpaired Student’s t-test (parametric comparison) or Mann-Whitney U-test (nonparametric comparison). For multiple comparisons, a one-way analysis of variance (ANOVA) (parametric) or Kruskal-Wallis test (nonparametric) with Tukey’s post hoc test were applied and adjusted *P* values are presented. *P* < 0.05 was considered statistically significant; exact *P* values are provided whenever possible in the figures and/or figure legends.

## Supplemental information

**Figure S1. Flow cytometry validates the heterogeneity and accumulation of CD8^+^ T lymphocyte subsets in the myelin mutant CNS** (**A**) Representative plots of flow cytometric analysis of single, viable CD45^high^CD8^+^ T cells freshly isolated from brains of 15-month-old *Wt* (top) and *PLPmut* (bottom) mice. Gated CD8^+^ T cells are analyzed for expression of CXCR4, CXCR6, Ly6A/E, and CD103. Percentages and identity of the respective cells are indicated in the quadrants. (**B**) CD8^+^ T cells comprise similar proportions of CXCR6^+^CXCR4^+^ cells (CAT5) and CXCR6^+^CXCR4^−^ cells (CAT1-4, IST) when comparing *Wt* and *PLPmut* mice (*n* = 6 mice per group). (**C**) Among CD8^+^CXCR6^+^CXCR4^−^ T cells, Ly6A/E^+^CD103^+^ cells (CAT2) but not Ly6A/E^+^CD103^−^ cells show an increased frequency. (**D**). Total numbers of CD45^high^CD8^+^ T cells and (**E**) the different subsets per brain reflect a significant accumulation of most populations with a disproportionally increased number of CAT2 cells. B-E: unpaired Student’s t-test. Data are presented as the mean ± SD. All data represent at least three independent experiments

**Figure S2. Immunohistochemistry validates the heterogeneity and activation of CD8^+^ T cell subsets in the myelin mutant CNS** (**A**) Immunofluorescence detection of CD8, LAG3, and CXCR4 or CD8, CD103, and Ly6A/E in the optic nerves of 9-month-old *Wt* and *PLPmut* mice. CD8^+^ T cells (arrows) show heterogenous expression of these markers. Scale bar, 10 μm. (**B**) Quantification of LAG3^+^ (CAT1), CXCR4^+^ (CAT5), CD103^+^ (CAT2) subsets among CD8+ T cells as well as Ly6A/E immunoreactivity of CAT2 (*n* = 5 mice per group). There is an increased frequency of CD103^+^ cells with increased Ly6A/E expression detectable among CD8^+^ T cells in *PLPmut* mice. B: unpaired Student’s t-test. Data are presented as the mean ± SD. All data represent at least three independent experiments.

**Figure S3. Sphingosine-1-receptor modulation with FTY720 depletes circulating T cells in *PLPmut* mice** (**A**) Flow cytometric quantification of leukocytes, CD11b^+^ myeloid cells, CD8^+^ T cells, and CD4^+^ T cells per μl blood from *Wt*, *PLPmut* and FTY720-treated *PLPmut* mice (*n* = 5 mice per group) after the preventive regimen shown in Fig. 3 B. (**B**) Representative immunohistochemical detection of CD8^+^ T cells in blood and spleen of *PLPmut* and FTY720-treated *PLPmut* mice. Scale bars, 20 μm (top) and 40 μm (bottom). Numbers of CD8^+^ T cells and spleen volume are strongly decreased by FY720. (**C**) Analysis of the relative spleen weights in *Wt*, *PLPmut* and FTY720-treated *PLPmut* mice (*n* = 5 mice per group) using regimens indicated in Fig. 3 B. Preventive FTY720 treatment reduces spleen weight in *PLPmut* mice which is restored after termination at half time. Therapeutic FTY720 treatment also reduces spleen weight. C: one-way ANOVA with Tukey’s multiple comparisons test. Data are presented as the mean ± SD. All data represent at least three independent experiments.

**Figure S4. Expression of Sn and CD11c identifies activated microglia in *PLPmut* mice** (**A**) Representative immunofluorescent detection of Sn on CD11b^+^ microglia in the optic nerves from 15-month-old *PLPmut* mice. Arrows indicate Sn^+^ cells. Quantifications are provided in Fig. 3 G. Scale bar, 20 μm. (**B**) Representative immunofluorescent detection of CD11c, Sn, and P2RY12 in the optic nerves from 15-month-old *PLPmut* mice. Arrows indicate CD11c^+^Sn^+^P2RY12^+^ cells. Sn and CD11c expression is restricted to activated microglia with reduced P2RY12 expression. Scale bar, 20 μm. All data represent at least three independent experiments.

**Figure S5. Beneficial effects of FTY720 treatment in *PLPmut* mice are immunomodulatory** (**A**) Quantification of SMI32^+^ axonal spheroids in the optic nerves, (**B**) RGCs, and (**G**) OCT analysis of the innermost retinal composite layer (NFL/GCL/IPL) in peripapillary circle scans in 9-month-old *PLPmut*/*Rag1^−/−^* mice with or without preventive FTY720 treatment (*n* = 5 mice per group). FTY720 treatment has no beneficial effect on the mild neurodegeneration observed in immunodeficient *PLPmut* mice. A-G: unpaired Student’s t-test. All data represent at least three independent experiments.

**Figure S6. Increased expression of MHC-I on oligodendrocytes and microglia in *PLPmut* mice** (**A**) Representative immunofluorescent detection of MHC-I on APC^+^ oligodendrocytes or (**B**) IBA1^+^ microglia in the optic nerves from 18-month-old *Wt* and *PLPmut* mice. Arrows indicate MHC-I^+^ cells. Quantifications are provided in Fig. 7 H and I. Scale bars, 20 μm. All data represent at least three independent experiments.

**Table S1.** Complete lists of cluster-specific marker and differentially expressed genes for the scRNA-seq data.

## References

1. Filley, C.M. (2021). White matter and human behavior. Science 372, 1265–1266. 10.1126/science.abj1881.

2. Salvadores, N., Sanhueza, M., Manque, P., and Court, F.A. (2017). Axonal Degeneration during Aging and Its Functional Role in Neurodegenerative Disorders. Front Neurosci 11, 451. 10.3389/fnins.2017.00451.

3. Stassart, R.M., Mobius, W., Nave, K.A., and Edgar, J.M. (2018). The Axon-Myelin Unit in Development and Degenerative Disease. Front Neurosci 12, 467. 10.3389/fnins.2018.00467.

4. Matejuk, A., Vandenbark, A.A., and Offner, H. (2021). Cross-Talk of the CNS With Immune Cells and Functions in Health and Disease. Front Neurol 12, 672455. 10.3389/fneur.2021.672455.

5. Kirby, L., and Castelo-Branco, G. (2021). Crossing boundaries: Interplay between the immune system and oligodendrocyte lineage cells. Semin Cell Dev Biol 116, 45–52. 10.1016/j.semcdb.2020.10.013.

6. Hughes, A.N. (2021). Glial Cells Promote Myelin Formation and Elimination. Front Cell Dev Biol 9, 661486. 10.3389/fcell.2021.661486.

7. Hickman, S., Izzy, S., Sen, P., Morsett, L., and El Khoury, J. (2018). Microglia in neurodegeneration. Nat Neurosci 21, 1359–1369. 10.1038/s41593-018-0242-x.

8. Schetters, S.T.T., Gomez-Nicola, D., Garcia-Vallejo, J.J., and Van Kooyk, Y. (2017). Neuroinflammation: Microglia and T Cells Get Ready to Tango. Front Immunol 8, 1905. 10.3389/fimmu.2017.01905.

9. Chitnis, T., and Weiner, H.L. (2017). CNS inflammation and neurodegeneration. J Clin Invest 127, 3577–3587. 10.1172/JCI90609.

10. Gleichman, A.J., and Carmichael, S.T. (2020). Glia in neurodegeneration: Drivers of disease or along for the ride? Neurobiol Dis 142, 104957. 10.1016/j.nbd.2020.104957.

11. Pandey, S., Shen, K., Lee, S.H., Shen, Y.A., Wang, Y., Otero-Garcia, M., Kotova, N., Vito, S.T., Laufer, B.I., Newton, D.F., et al. (2022). Disease-associated oligodendrocyte responses across neurodegenerative diseases. Cell Rep 40, 111189. 10.1016/j.celrep.2022.111189.

12. Carrasco, E., Gomez de Las Heras, M.M., Gabande-Rodriguez, E., Desdin-Mico, G., Aranda, J.F., and Mittelbrunn, M. (2021). The role of T cells in age-related diseases. Nat Rev Immunol. 10.1038/s41577-021-00557-4.

13. Galiano-Landeira, J., Torra, A., Vila, M., and Bove, J. (2020). CD8 T cell nigral infiltration precedes synucleinopathy in early stages of Parkinson’s disease. Brain 143, 3717–3733. 10.1093/brain/awaa269.

14. Gate, D., Saligrama, N., Leventhal, O., Yang, A.C., Unger, M.S., Middeldorp, J., Chen, K., Lehallier, B., Channappa, D., De Los Santos, M.B., et al. (2020). Clonally expanded CD8 T cells patrol the cerebrospinal fluid in Alzheimer’s disease. Nature 577, 399–404. 10.1038/s41586-019-1895-7.

15. Groh, J., and Martini, R. (2017). Neuroinflammation as modifier of genetically caused neurological disorders of the central nervous system: Understanding pathogenesis and chances for treatment. Glia 65, 1407–1422. 10.1002/glia.23162.

16. Groh, J., Friedman, H.C., Orel, N., Ip, C.W., Fischer, S., Spahn, I., Schaffner, E., Horner, M., Stadler, D., Buttmann, M., et al. (2016). Pathogenic inflammation in the CNS of mice carrying human PLP1 mutations. Hum Mol Genet 25, 4686–4702. 10.1093/hmg/ddw296.

17. Ip, C.W., Kroner, A., Bendszus, M., Leder, C., Kobsar, I., Fischer, S., Wiendl, H., Nave, K.A., and Martini, R. (2006). Immune cells contribute to myelin degeneration and axonopathic changes in mice overexpressing proteolipid protein in oligodendrocytes. J Neurosci 26, 8206–8216. 10.1523/JNEUROSCI.1921-06.2006.

18. Ip, C.W., Kroner, A., Groh, J., Huber, M., Klein, D., Spahn, I., Diem, R., Williams, S.K., Nave, K.A., Edgar, J.M., and Martini, R. (2012). Neuroinflammation by cytotoxic T-lymphocytes impairs retrograde axonal transport in an oligodendrocyte mutant mouse. PLoS One 7, e42554. 10.1371/journal.pone.0042554.

19. Kroner, A., Ip, C.W., Thalhammer, J., Nave, K.A., and Martini, R. (2010). Ectopic T-cell specificity and absence of perforin and granzyme B alleviate neural damage in oligodendrocyte mutant mice. Am J Pathol 176, 549–555. 10.2353/ajpath.2010.090722.

20. Groh, J., Klein, D., Berve, K., West, B.L., and Martini, R. (2019). Targeting microglia attenuates neuroinflammation-related neural damage in mice carrying human PLP1 mutations. Glia 67, 277–290. 10.1002/glia.23539.

21. Groh, J., Horner, M., and Martini, R. (2018). Teriflunomide attenuates neuroinflammation-related neural damage in mice carrying human PLP1 mutations. J Neuroinflammation 15, 194. 10.1186/s12974-018-1228-z.

22. Factor, D.C., Barbeau, A.M., Allan, K.C., Hu, L.R., Madhavan, M., Hoang, A.T., Hazel, K.E.A., Hall, P.A., Nisraiyya, S., Najm, F.J., et al. (2020). Cell Type-Specific Intralocus Interactions Reveal Oligodendrocyte Mechanisms in MS. Cell 181, 382–395 e321. 10.1016/j.cell.2020.03.002.

23. Caprariello, A.V., Rogers, J.A., Morgan, M.L., Hoghooghi, V., Plemel, J.R., Koebel, A., Tsutsui, S., Dunn, J.F., Kotra, L.P., Ousman, S.S., et al. (2018). Biochemically altered myelin triggers autoimmune demyelination. Proc Natl Acad Sci U S A 115, 5528–5533. 10.1073/pnas.1721115115.

24. Meijer, M., Agirre, E., Kabbe, M., van Tuijn, C.A., Heskol, A., Zheng, C., Mendanha Falcao, A., Bartosovic, M., Kirby, L., Calini, D., et al. (2022). Epigenomic priming of immune genes implicates oligodendroglia in multiple sclerosis susceptibility. Neuron 110, 1193–1210 e1113. 10.1016/j.neuron.2021.12.034.

25. Groh, J., Knöpper, K., Arampatzi, P., Yuan, X., Lößlein, L., Saliba, A.-E., Kastenmüller, W., and Martini, R. (2021). Accumulation of cytotoxic T cells in the aged CNS leads to axon degeneration and contributes to cognitive and motor decline. Nature Aging 1, 357–367. 10.1038/s43587-021-00049-z.

26. McGinley, M.P., and Cohen, J.A. (2021). Sphingosine 1-phosphate receptor modulators in multiple sclerosis and other conditions. Lancet 398, 1184–1194. 10.1016/S0140-6736(21)00244-0.

27. Szabo, P.A., Levitin, H.M., Miron, M., Snyder, M.E., Senda, T., Yuan, J., Cheng, Y.L., Bush, E.C., Dogra, P., Thapa, P., et al. (2019). Single-cell transcriptomics of human T cells reveals tissue and activation signatures in health and disease. Nat Commun 10, 4706. 10.1038/s41467-019-12464-3.

28. Khan, A.B., Carpenter, B., Santos, E.S.P., Pospori, C., Khorshed, R., Griffin, J., Velica, P., Zech, M., Ghorashian, S., Forrest, C., et al. (2018). Redirection to the bone marrow improves T cell persistence and antitumor functions. J Clin Invest 128, 2010–2024. 10.1172/JCI97454.

29. Schirmer, L., Velmeshev, D., Holmqvist, S., Kaufmann, M., Werneburg, S., Jung, D., Vistnes, S., Stockley, J.H., Young, A., Steindel, M., et al. (2019). Neuronal vulnerability and multilineage diversity in multiple sclerosis. Nature 573, 75–82. 10.1038/s41586-019-1404-z.

30. Falcao, A.M., van Bruggen, D., Marques, S., Meijer, M., Jakel, S., Agirre, E., Samudyata, Floriddia, E.M., Vanichkina, D.P., Ffrench-Constant, C., et al. (2018). Disease-specific oligodendrocyte lineage cells arise in multiple sclerosis. Nat Med 24, 1837–1844. 10.1038/s41591-018-0236-y.

31. Kirby, L., Jin, J., Cardona, J.G., Smith, M.D., Martin, K.A., Wang, J., Strasburger, H., Herbst, L., Alexis, M., Karnell, J., et al. (2019). Oligodendrocyte precursor cells present antigen and are cytotoxic targets in inflammatory demyelination. Nat Commun 10, 3887. 10.1038/s41467-019-11638-3.

32. Kaya, T., Mattugini, N., Liu, L., Ji, H., Cantuti-Castelvetri, L., Wu, J., Schifferer, M., Groh, J., Martini, R., Besson-Girard, S., et al. (2022). CD8(+) T cells induce interferon-responsive oligodendrocytes and microglia in white matter aging. Nat Neurosci 25, 1446–1457. 10.1038/s41593-022-01183-6.

33. Stys, P.K., Zamponi, G.W., van Minnen, J., and Geurts, J.J. (2012). Will the real multiple sclerosis please stand up? Nat Rev Neurosci 13, 507–514. 10.1038/nrn3275.

34. Trapp, B.D., and Nave, K.A. (2008). Multiple sclerosis: an immune or neurodegenerative disorder? Annu Rev Neurosci 31, 247–269. 10.1146/annurev.neuro.30.051606.094313.

35. Luchicchi, A., Preziosa, P., and t Hart, B. (2021). Editorial: “Inside-Out” vs “Outside-In” Paradigms in Multiple Sclerosis Etiopathogenesis. Front Cell Neurosci 15, 666529. 10.3389/fncel.2021.666529.

36. Kenigsbuch, M., Bost, P., Halevi, S., Chang, Y., Chen, S., Ma, Q., Hajbi, R., Schwikowski, B., Bodenmiller, B., Fu, H., et al. (2022). A shared disease-associated oligodendrocyte signature among multiple CNS pathologies. Nat Neurosci 25, 876–886. 10.1038/s41593-022-01104-7.

37. Wakim, L.M., Woodward-Davis, A., Liu, R., Hu, Y., Villadangos, J., Smyth, G., and Bevan, M.J. (2012). The molecular signature of tissue resident memory CD8 T cells isolated from the brain. J Immunol 189, 3462–3471. 10.4049/jimmunol.1201305.

38. Smolders, J., Heutinck, K.M., Fransen, N.L., Remmerswaal, E.B.M., Hombrink, P., Ten Berge, I.J.M., van Lier, R.A.W., Huitinga, I., and Hamann, J. (2018). Tissue-resident memory T cells populate the human brain. Nat Commun 9, 4593. 10.1038/s41467-018-07053-9.

39. Urban, S.L., Jensen, I.J., Shan, Q., Pewe, L.L., Xue, H.H., Badovinac, V.P., and Harty, J.T. (2020). Peripherally induced brain tissue-resident memory CD8(+) T cells mediate protection against CNS infection. Nat Immunol 21, 938–949. 10.1038/s41590-020-0711-8.

40. Schenkel, J.M., and Masopust, D. (2014). Tissue-resident memory T cells. Immunity 41, 886–897. 10.1016/j.immuni.2014.12.007.

41. Evrard, M., Wynne-Jones, E., Peng, C., Kato, Y., Christo, S.N., Fonseca, R., Park, S.L., Burn, T.N., Osman, M., Devi, S., et al. (2022). Sphingosine 1-phosphate receptor 5 (S1PR5) regulates the peripheral retention of tissue-resident lymphocytes. J Exp Med 219. 10.1084/jem.20210116.

42. Frieser, D., Pignata, A., Khajavi, L., Shlesinger, D., Gonzalez-Fierro, C., Nguyen, X.H., Yermanos, A., Merkler, D., Hoftberger, R., Desestret, V., et al. (2022). Tissue-resident CD8(+) T cells drive compartmentalized and chronic autoimmune damage against CNS neurons. Sci Transl Med 14, eabl6157. 10.1126/scitranslmed.abl6157.

43. Vincenti, I., Page, N., Steinbach, K., Yermanos, A., Lemeille, S., Nunez, N., Kreutzfeldt, M., Klimek, B., Di Liberto, G., Egervari, K., et al. (2022). Tissue-resident memory CD8(+) T cells cooperate with CD4(+) T cells to drive compartmentalized immunopathology in the CNS. Sci Transl Med 14, eabl6058. 10.1126/scitranslmed.abl6058.

44. Machado-Santos, J., Saji, E., Troscher, A.R., Paunovic, M., Liblau, R., Gabriely, G., Bien, C.G., Bauer, J., and Lassmann, H. (2018). The compartmentalized inflammatory response in the multiple sclerosis brain is composed of tissue-resident CD8+ T lymphocytes and B cells. Brain 141, 2066–2082. 10.1093/brain/awy151.

45. Beltran, E., Gerdes, L.A., Hansen, J., Flierl-Hecht, A., Krebs, S., Blum, H., Ertl-Wagner, B., Barkhof, F., Kumpfel, T., Hohlfeld, R., and Dornmair, K. (2019). Early adaptive immune activation detected in monozygotic twins with prodromal multiple sclerosis. J Clin Invest 129, 4758–4768. 10.1172/JCI128475.

46. Lassmann, H. (2018). Pathogenic Mechanisms Associated With Different Clinical Courses of Multiple Sclerosis. Front Immunol 9, 3116. 10.3389/fimmu.2018.03116.

47. Smolders, J., van Luijn, M.M., Hsiao, C.C., and Hamann, J. (2022). T-cell surveillance of the human brain in health and multiple sclerosis. Semin Immunopathol. 10.1007/s00281-022-00926-8.

48. Giovannoni, G., Popescu, V., Wuerfel, J., Hellwig, K., Iacobeus, E., Jensen, M.B., Garcia-Dominguez, J.M., Sousa, L., De Rossi, N., Hupperts, R., et al. (2022). Smouldering multiple sclerosis: the ‘real MS’. Ther Adv Neurol Disord 15, 17562864211066751. 10.1177/17562864211066751.

49. Zalocusky, K.A., Najm, R., Taubes, A.L., Hao, Y., Yoon, S.Y., Koutsodendris, N., Nelson, M.R., Rao, A., Bennett, D.A., Bant, J., et al. (2021). Neuronal ApoE upregulates MHC-I expression to drive selective neurodegeneration in Alzheimer’s disease. Nat Neurosci 24, 786–798. 10.1038/s41593-021-00851-3.

50. Standiford, M.M., Grund, E.M., and Howe, C.L. (2021). Citrullinated myelin induces microglial TNFalpha and inhibits endogenous repair in the cuprizone model of demyelination. J Neuroinflammation 18, 305. 10.1186/s12974-021-02360-3.

51. Funfschilling, U., Supplie, L.M., Mahad, D., Boretius, S., Saab, A.S., Edgar, J., Brinkmann, B.G., Kassmann, C.M., Tzvetanova, I.D., Mobius, W., et al. (2012). Glycolytic oligodendrocytes maintain myelin and long-term axonal integrity. Nature 485, 517–521. 10.1038/nature11007.

52. Sobottka, B., Harrer, M.D., Ziegler, U., Fischer, K., Wiendl, H., Hunig, T., Becher, B., and Goebels, N. (2009). Collateral bystander damage by myelin-directed CD8+ T cells causes axonal loss. Am J Pathol 175, 1160–1166. 10.2353/ajpath.2009.090340.

53. Witte, M.E., Schumacher, A.M., Mahler, C.F., Bewersdorf, J.P., Lehmitz, J., Scheiter, A., Sanchez, P., Williams, P.R., Griesbeck, O., Naumann, R., et al. (2019). Calcium Influx through Plasma-Membrane Nanoruptures Drives Axon Degeneration in a Model of Multiple Sclerosis. Neuron 101, 615–624 e615. 10.1016/j.neuron.2018.12.023.

54. Schäffner, E., Edgar, J., Lehning, M., Strauß, J., Bosch-Queralt, M., Wieghofer, P., Berghoff, S., Krueger, M., Morawski, M., Reinert, T., et al. (2021). Myelin insulation as a risk factor for axonal degeneration in autoimmune demyelinating disease. bioRxiv, 2021.2011.2011.468223. 10.1101/2021.11.11.468223.

55. Groh, J., Berve, K., and Martini, R. (2021). Immune modulation attenuates infantile neuronal ceroid lipofuscinosis in mice before and after disease onset. Brain Commun 3, fcab047. 10.1093/braincomms/fcab047.

56. Lublin, F., Miller, D.H., Freedman, M.S., Cree, B.A.C., Wolinsky, J.S., Weiner, H., Lubetzki, C., Hartung, H.P., Montalban, X., Uitdehaag, B.M.J., et al. (2016). Oral fingolimod in primary progressive multiple sclerosis (INFORMS): a phase 3, randomised, double-blind, placebo-controlled trial. Lancet 387, 1075–1084. 10.1016/S0140-6736(15)01314-8.

57. Tilly, G., Cadoux, M., Garcia, A., Morille, J., Wiertlewski, S., Pecqueur, C., Brouard, S., Laplaud, D., and Degauque, N. (2021). Teriflunomide Treatment of Multiple Sclerosis Selectively Modulates CD8 Memory T Cells. Front Immunol 12, 730342. 10.3389/fimmu.2021.730342.

58. Medina, S., Sainz de la Maza, S., Villarrubia, N., Alvarez-Lafuente, R., Costa-Frossard, L., Arroyo, R., Monreal, E., Tejeda-Velarde, A., Rodriguez-Martin, E., Roldan, E., et al. (2019). Teriflunomide induces a tolerogenic bias in blood immune cells of MS patients. Ann Clin Transl Neurol 6, 355–363. 10.1002/acn3.711.

59. Akane, K., Kojima, S., Mak, T.W., Shiku, H., and Suzuki, H. (2016). CD8+CD122+CD49dlow regulatory T cells maintain T-cell homeostasis by killing activated T cells via Fas/FasL-mediated cytotoxicity. Proc Natl Acad Sci U S A 113, 2460–2465. 10.1073/pnas.1525098113.

60. Mishra, S., Srinivasan, S., Ma, C., and Zhang, N. (2021). CD8(+) Regulatory T Cell - A Mystery to Be Revealed. Front Immunol 12, 708874. 10.3389/fimmu.2021.708874.

61. Dai, H., Wan, N., Zhang, S., Moore, Y., Wan, F., and Dai, Z. (2010). Cutting edge: programmed death-1 defines CD8+CD122+ T cells as regulatory versus memory T cells. J Immunol 185, 803–807. 10.4049/jimmunol.1000661.

62. Saligrama, N., Zhao, F., Sikora, M.J., Serratelli, W.S., Fernandes, R.A., Louis, D.M., Yao, W., Ji, X., Idoyaga, J., Mahajan, V.B., et al. (2019). Opposing T cell responses in experimental autoimmune encephalomyelitis. Nature 572, 481–487. 10.1038/s41586-019-1467-x.

63. Groh, J., Ribechini, E., Stadler, D., Schilling, T., Lutz, M.B., and Martini, R. (2016). Sialoadhesin promotes neuroinflammation-related disease progression in two mouse models of CLN disease. Glia 64, 792–809. 10.1002/glia.22962.

64. Li, J., Zaslavsky, M., Su, Y., Guo, J., Sikora, M.J., van Unen, V., Christophersen, A., Chiou, S.H., Chen, L., Li, J., et al. (2022). KIR(+)CD8(+) T cells suppress pathogenic T cells and are active in autoimmune diseases and COVID-19. Science, eabi9591. 10.1126/science.abi9591.

65. Yu, F.X., Chai, T.F., He, H., Hagen, T., and Luo, Y. (2010). Thioredoxin-interacting protein (Txnip) gene expression: sensing oxidative phosphorylation status and glycolytic rate. J Biol Chem 285, 25822–25830. 10.1074/jbc.M110.108290.

66. Mombaerts, P., Iacomini, J., Johnson, R.S., Herrup, K., Tonegawa, S., and Papaioannou, V.E. (1992). RAG-1-deficient mice have no mature B and T lymphocytes. Cell 68, 869–877. 10.1016/0092-8674(92)90030-g.

67. Heusel, J.W., Wesselschmidt, R.L., Shresta, S., Russell, J.H., and Ley, T.J. (1994). Cytotoxic lymphocytes require granzyme B for the rapid induction of DNA fragmentation and apoptosis in allogeneic target cells. Cell 76, 977–987. 10.1016/0092-8674(94)90376-x.

68. Hogquist, K.A., Jameson, S.C., Heath, W.R., Howard, J.L., Bevan, M.J., and Carbone, F.R. (1994). T cell receptor antagonist peptides induce positive selection. Cell 76, 17–27. 10.1016/0092-8674(94)90169-4.

69. Zheng, G.X., Terry, J.M., Belgrader, P., Ryvkin, P., Bent, Z.W., Wilson, R., Ziraldo, S.B., Wheeler, T.D., McDermott, G.P., Zhu, J., et al. (2017). Massively parallel digital transcriptional profiling of single cells. Nat Commun 8, 14049. 10.1038/ncomms14049.

70. Dobin, A., Davis, C.A., Schlesinger, F., Drenkow, J., Zaleski, C., Jha, S., Batut, P., Chaisson, M., and Gingeras, T.R. (2013). STAR: ultrafast universal RNA-seq aligner. Bioinformatics 29, 15–21. 10.1093/bioinformatics/bts635.

71. Satija, R., Farrell, J.A., Gennert, D., Schier, A.F., and Regev, A. (2015). Spatial reconstruction of single-cell gene expression data. Nat Biotechnol 33, 495–502. 10.1038/nbt.3192.

72. Groh, J., Berve, K., and Martini, R. (2017). Fingolimod and Teriflunomide Attenuate Neurodegeneration in Mouse Models of Neuronal Ceroid Lipofuscinosis. Mol Ther 25, 1889–1899. 10.1016/j.ymthe.2017.04.021.

73. Metzler, B., Gfeller, P., Wieczorek, G., Li, J., Nuesslein-Hildesheim, B., Katopodis, A., Mueller, M., and Brinkmann, V. (2008). Modulation of T cell homeostasis and alloreactivity under continuous FTY720 exposure. Int Immunol 20, 633–644. 10.1093/intimm/dxn023.

74. Nair, A.B., and Jacob, S. (2016). A simple practice guide for dose conversion between animals and human. J Basic Clin Pharm 7, 27–31. 10.4103/0976-0105.177703.

75. Groh, J., Stadler, D., Buttmann, M., and Martini, R. (2014). Non-invasive assessment of retinal alterations in mouse models of infantile and juvenile neuronal ceroid lipofuscinosis by spectral domain optical coherence tomography. Acta Neuropathol Commun 2, 54. 10.1186/2051-5960-2-54.

76. Groh, J., Kuhl, T.G., Ip, C.W., Nelvagal, H.R., Sri, S., Duckett, S., Mirza, M., Langmann, T., Cooper, J.D., and Martini, R. (2013). Immune cells perturb axons and impair neuronal survival in a mouse model of infantile neuronal ceroid lipofuscinosis. Brain 136, 1083–1101. 10.1093/brain/awt020.

77. Faul, F., Erdfelder, E., Lang, A.G., and Buchner, A. (2007). G*Power 3: a flexible statistical power analysis program for the social, behavioral, and biomedical sciences. Behav Res Methods 39, 175–191. 10.3758/bf03193146.

