## Supplementary figures and images for "Cytotoxic CNS-associated T cells drive axon degeneration by targeting perturbed myelinating oligodendrocytes in *PLP1* mutant mice"

### Figures S1-S6

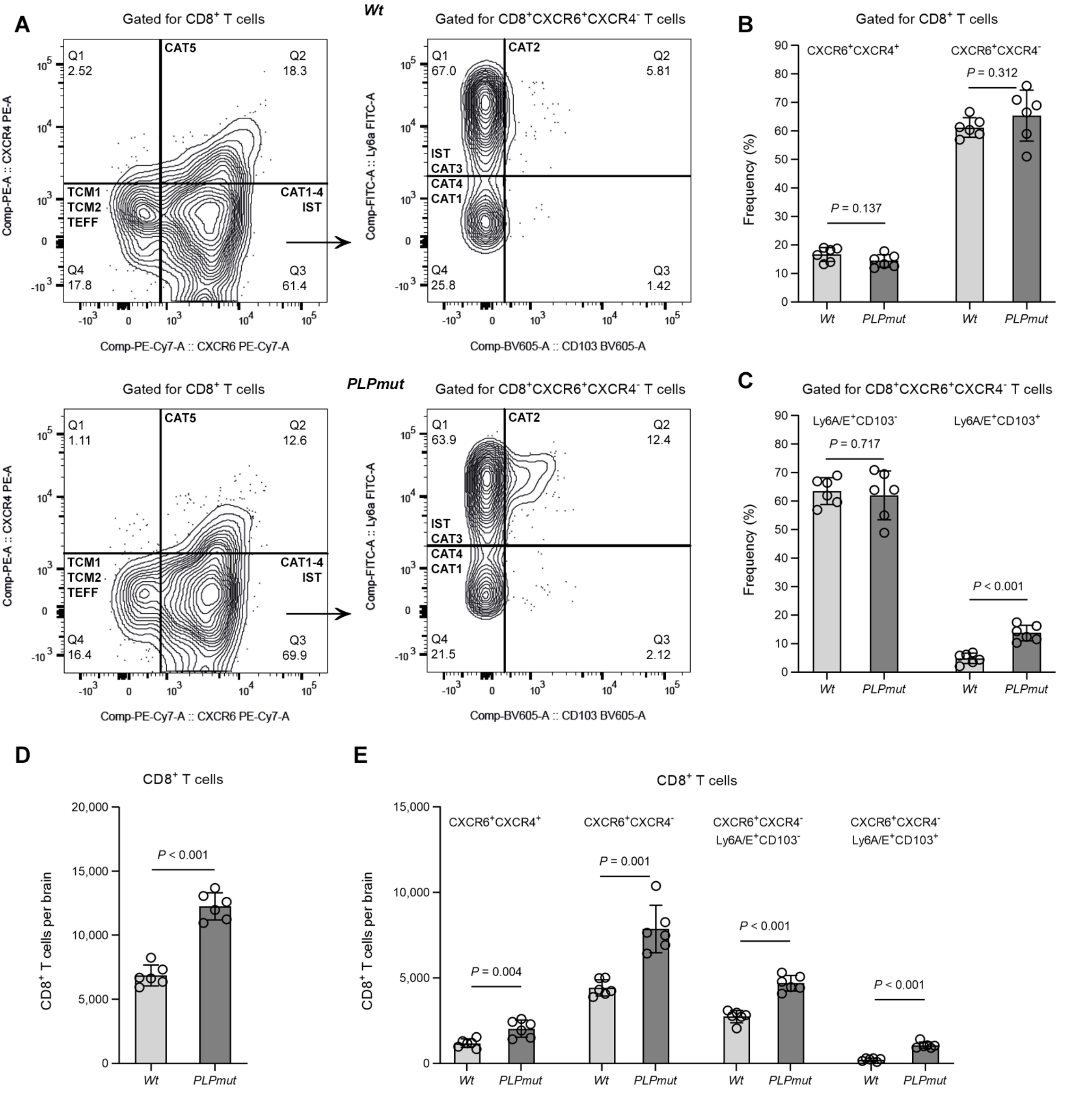

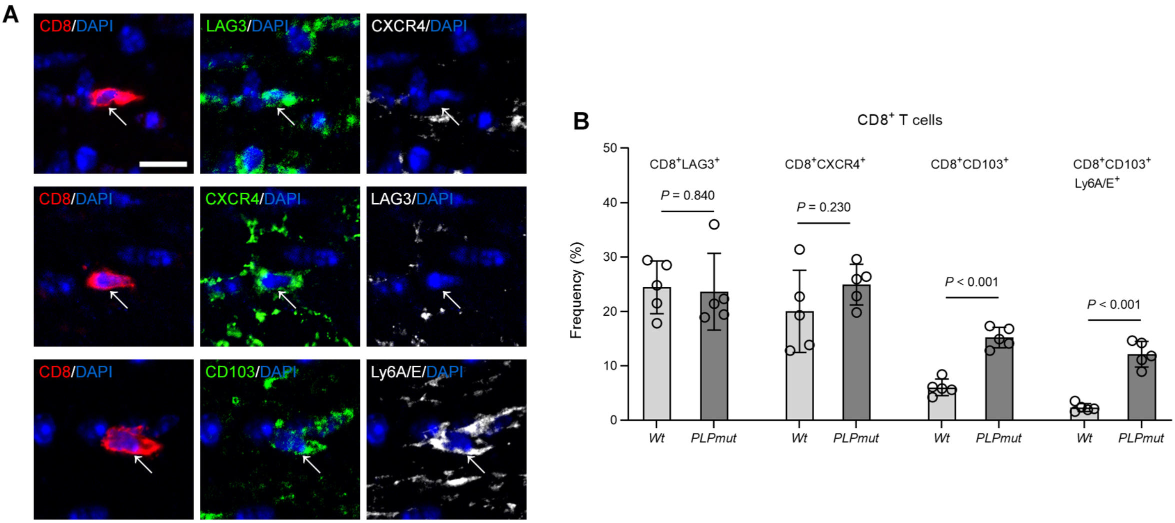

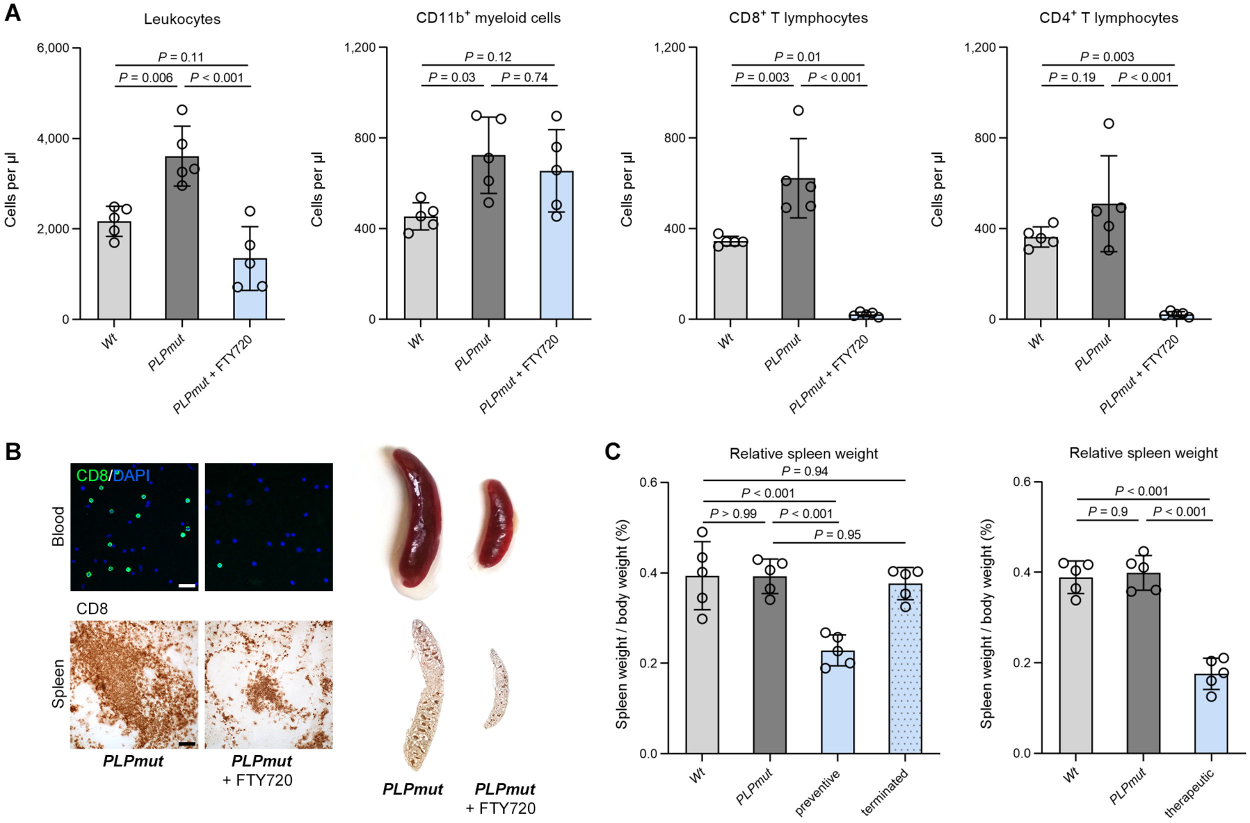

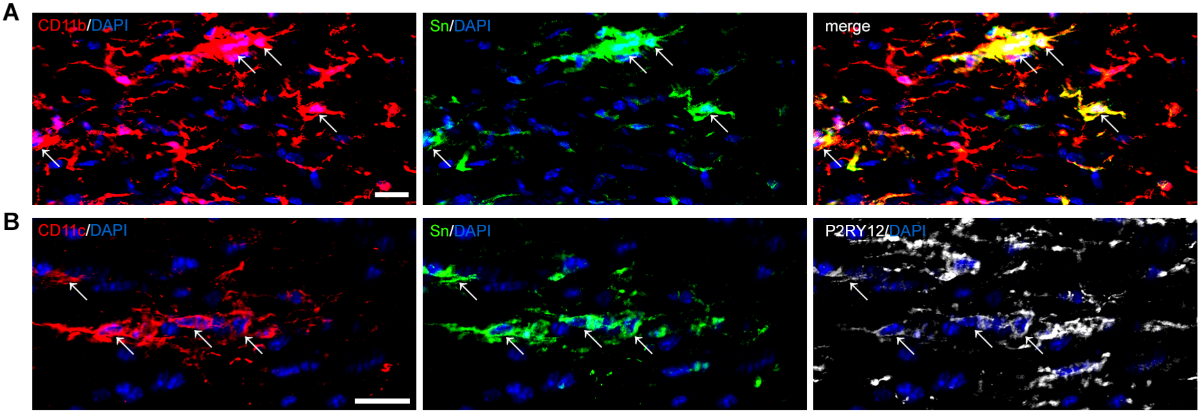


**Figure S4**

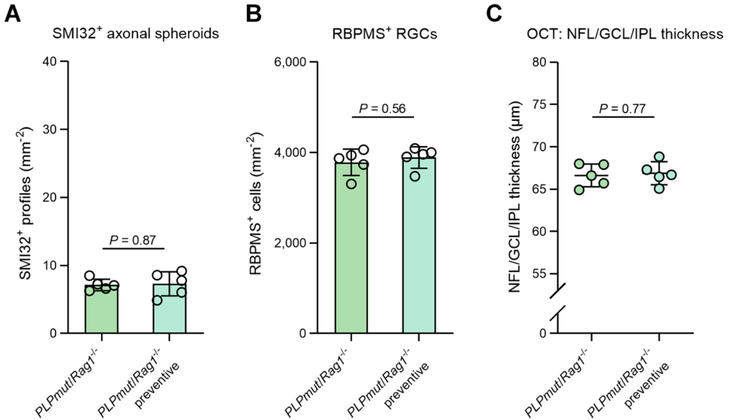

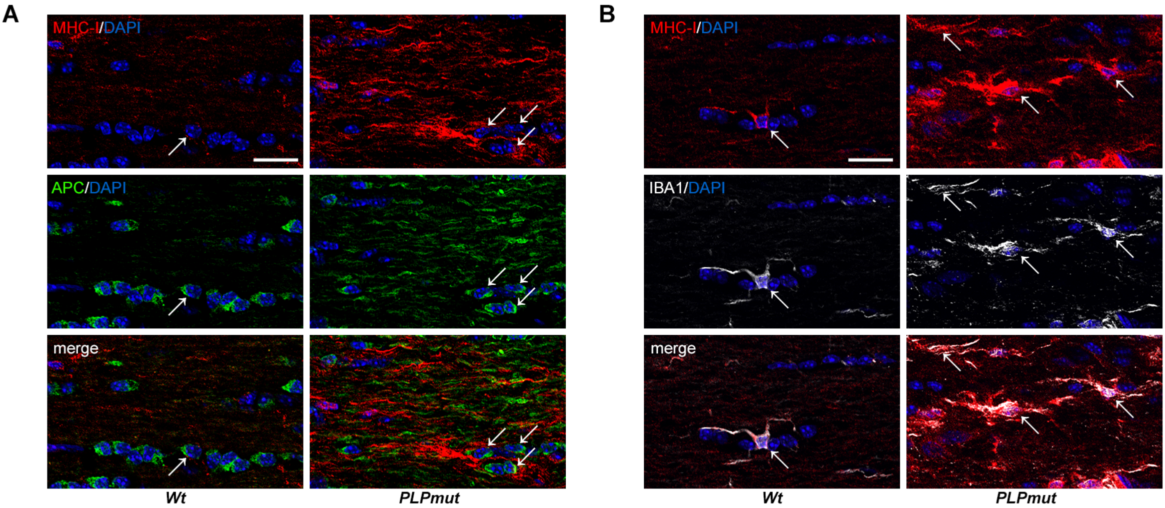
